# The regional distribution of resident immune cells shapes distinct immunological environments along the murine epididymis

**DOI:** 10.1101/2022.06.13.495924

**Authors:** Christiane Pleuger, Dingding Ai, Minea Hoppe, Laura Winter, Daniel Bohnert, Dominik Karl, Stefan Guenther, Slava Epelman, Crystal Kantores, Monika Fijak, Sarina Ravens, Ralf Middendorff, Johannes U. Mayer, Kate L. Loveland, Mark P. Hedger, Sudhanshu Bhushan, Andreas Meinhardt

## Abstract

The epididymis constitutes an important transition zone for post-testicular sperm maturation and storage. As the organ consists of a single convoluted duct, inflammation-associated tissue damage has a severe impact on fertility. In order to clarify the reasons for region-specific differences in the intensity of immune responses observed in a mouse model of acute bacterial epididymitis, we investigated the heterogeneity of resident immune cell populations within the epididymis under physiological conditions by scRNASeq analysis of extravascular CD45^+^ cells. 12 distinct immune cell subsets were identified, displaying substantial differences in distribution along the epididymis. Several distinct subsets of macrophages constituted the majority of these cells. Crucially, the proximal and distal regions showed striking differences in their immunological landscapes. These findings indicate that resident immune cells are strategically positioned along the epididymal duct, potentially providing different immunological environments required for sperm maturations and elimination of pathogens ascending the urogenital tract.

## Introduction

Within the male reproductive tract, the epididymis plays an essential role in post-testicular sperm maturation and storage. Immotile spermatozoa released from the seminiferous epithelium of the testis enter the epididymis via the efferent ducts and undergo distinct consecutive biochemical maturation processes required to gain motility and fertilization capacities (Belleannee et al., 2011; Skerget et al., 2015; Björkgren and Sipilä, 2019; Barrachina et al., 2022). The sequential maturation process is orchestrated by the pseudostratified epithelium composed of several different epithelial (principal, basal, narrow/clear) and immune cell types that creates an unique luminal milieu. The barrier function of the epididymal epithelium highly depends on epithelial integrity (Breton et al., 2019). Intraepithelial immune cells, particularly mononuclear phagocytes (MP), are highly abundant within the epididymal epithelium and perform a key role in the preservation of epithelial integrity (Smith et al., 2014).

From an immunological perspective, the epididymis performs a functionally complex role by providing an immunotolerant environment for transiting immunogenic spermatozoa, while maintaining the capacity to effectively combat invading pathogens ascending from the urethra and vas deferens. Previous investigations in rodents revealed differences in the immune reactions of opposing ends of the epididymis toward ascending bacterial infection and other inflammatory stimuli. In this regard, the proximal regions appear to be almost unresponsive, while the distal regions are prone to intense immune responses resulting in persistent tissue damage (Michel et al., 2016; Silva et al., 2018; Klein et al., 2019; Wang et al., 2019; Klein et al., 2020; Wijayarathna et al., 2020).

As the epididymis consists of a single highly convoluted duct that meanders through structurally different regions (initial segment [IS], caput [CT], corpus [CS], cauda [CD]), inflammation-associated tissue damage and fibrotic remodeling result in epididymal duct obstruction which has a direct impact on the maturation and passage of sperm and, thereby, fertility. The histopathological observations in rodent models replicate many of the clinical manifestations in epididymitis patients (Pilatz et al., 2015; Fijak et al., 2018). Epididymitis in humans is mostly caused by urogenital tract infections with coliform bacteria (i.e. uropathogenic *Escherichia coli* (UPEC)) or pathogens linked to sexually transmitted diseases (e.g. *Chlamydia trachomatis*, (Pilatz et al., 2015; Pleuger et al., 2020)) and can effectively be treated with antibiotics. However, up to 40% of epididymitis patients exhibit a persistent sub- or infertility (Rusz et al., 2012), most likely due to epididymal duct stenosis/obstruction and concomitant oligo- or azoospermia. The reasons underlying differences in different immune responsiveness, with strong pro-inflammatory immune response largely confined to the cauda, are not well-understood.

Within the last few decades, initial steps have been made in characterizing the immunological landscape within the epididymis and understanding how the epididymis is prepared for its immunological challenges (Nashan et al., 1989; Flickinger et al., 1997; Serre and Robaire, 1999; Da Silva et al., 2011; Shum et al., 2014; Pierucci-Alves et al., 2018; Voisin et al., 2018; Battistone et al., 2020; Mendelsohn et al., 2020; Wang et al., 2021). The murine epididymis is populated by various myeloid and lymphoid cell populations that are differentially distributed along the epididymal duct. Subsets of the MP system are the most prominent group within the epididymis and form a dense network within and around the epididymal duct, especially within the IS which is the site of spermatozoa entry (Da Silva et al., 2011; Battistone et al., 2020). Generally, the MP system comprises multiple subsets that can share similar cell surface markers, yet possess distinct functions related to tissue homeostasis and pathogen-specific immunity. Despite accumulating information about the localization and antigen-presentation and –processing properties (Da Silva et al., 2011; Da Silva and Smith, 2015; Battistone et al., 2020; Mendelsohn et al., 2020), the identity of MP subgroups within the epididymis as well as the full extent of their heterogeneity is still not well-understood mainly due to the general similarities between macrophage and DC subpopulations. In view of the fundamentally different immunological requirements of the epididymis, maintaining both a stable and immunotolerant microenvironment for sperm maturation in the proximal regions and the ability to mount adequate immune responses towards invading bacteria at the distal end, detailed investigation of the phenotypes, localization and function of resident immune cells is essential.

In this regard, we hypothesized that strategically positioned resident immune cells that function as both ‘scavengers’ and ‘guardians’ create distinct immunological landscapes within epididymal regions. These, in turn, are responsible for the observed differences in the intensity of the immune responses toward infectious or inflammatory stimuli as well as for tissue homeostasis and the maintenance of epithelial function that is essential for regulating the sequential steps of sperm maturation. Therefore, in this study, we aimed to both (i) analyze the differential immune responses to UPEC-elicited epididymitis, and (ii) uncover the immune diversity among epididymal regions, by using an unbiased single-cell RNA sequencing (scRNASeq) analysis complemented by flow cytometry and immunofluorescence analysis to localize identified populations *in situ*.

## Results

### Caput and cauda epididymides react fundamentally differently during acute bacterial epididymitis

To better understand the different immune responses within the epididymal regions and to expand on our previous studies, an experimental bacterial mouse epididymitis model was used to monitor disease progression up to 10 days post-infection (p.i., Fig. 1A). Bacteria were found in all epididymal regions (IS, caput, corpus, cauda) and in the testis one day p.i. (Fig. 1B), but persisted at high numbers only for up to 10 days in the cauda (Fig. 1B). Later time points were not examined, as it is known that bacteria are cleared towards day 30 p.i. (Klein et al., 2019). The caput showed no gross morphological alterations (Fig. 1C, Suppl. Fig. S1A-B), although slight histopathological changes, including mild focal epithelial damage and minor connective tissue deposition within the interstitium, were observed 5 days p.i. (Fig. 1C) resulting in an elevated disease score that returned to normal values at 10 days p.i. (Fig. 1D). In accordance with previous reports (Klein et al., 2020) severe tissue remodeling was seen in the cauda (Fig. 1E, Suppl. Fig. S1A-E) characterized by infiltration of immune cells, loss of epithelial integrity, connective tissue deposition in the interstitium, reduction of luminal diameter and ultimately, epididymal duct destruction resulting in a significantly increased and persistent overall disease score (Fig. 1F). Initially, immune cell infiltrates were predominantly located peripherally within the cauda (5 days p.i.) before larger leukocytic conglomerates/granulomas developed within the entire cauda region (Suppl. Fig. S1E). Sham control mice initially showed histopathological alterations in the cauda that were milder than in infected animals and returned to a level comparable to untreated epididymis towards day 10 p.i. (Fig. 1E, Fig. 1F, Suppl. Fig. S1E).

**Figure 1:**
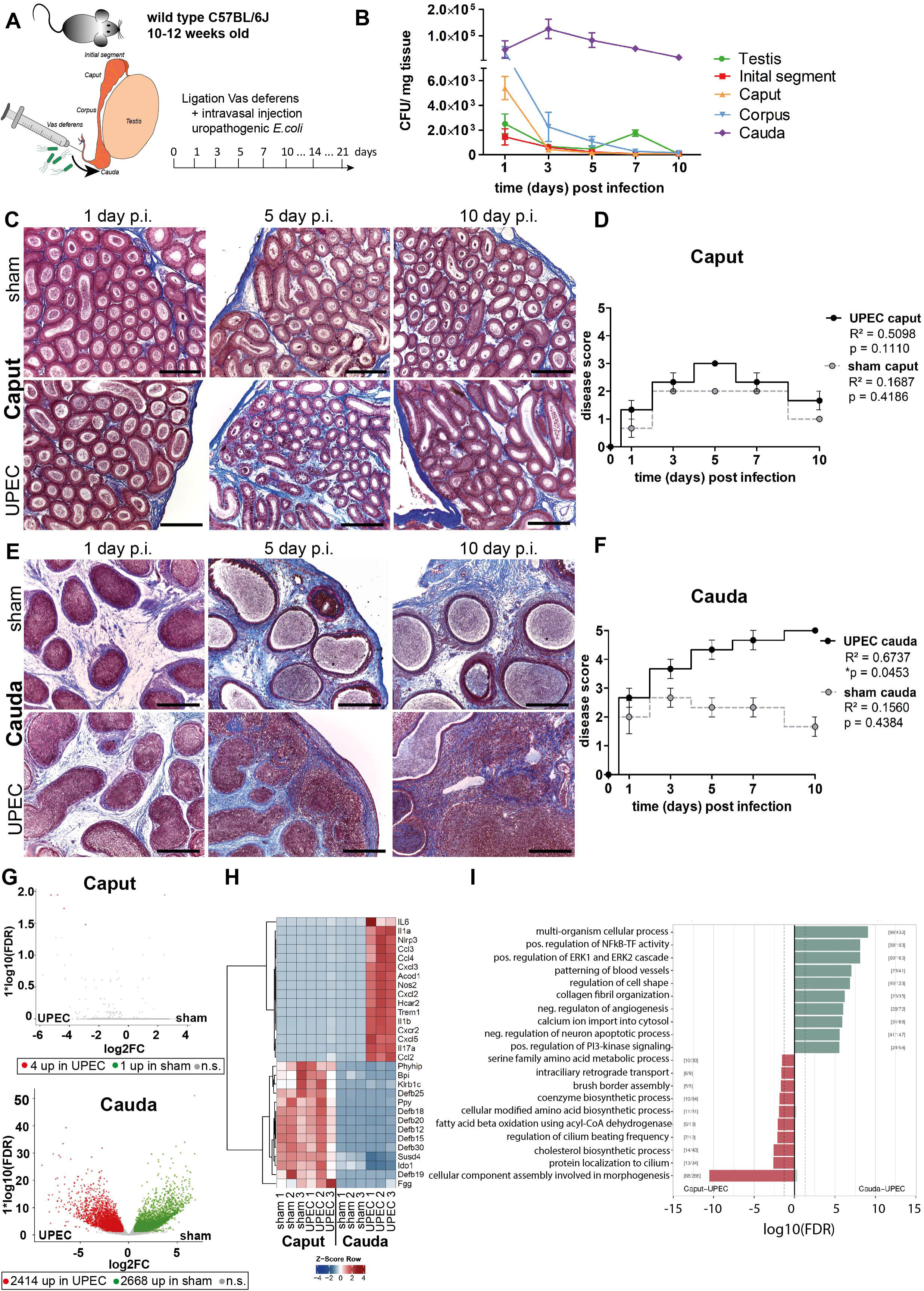
Analysis of differential immune responses of caput and cauda epididymides following UPEC infection in C57BL/6J wild type mice. **(A)** Male C57BL/6J mice (10-12 weeks old) were intravasally injected with UPEC or saline vehicle alone (sham) after ligation of the vas deferens. For the study organs were harvested and analyzed at the indicated time points. **(B)** Bacterial loads were assessed by determining colony forming units per mg tissue at the indicated time points within testis and the four main epididymal regions (IS, caput, corpus, cauda; n=4 per time point, mean ± SD). **(C and D)** Modified Masson-Goldner trichrome staining of caput (C) and cauda (D) epididymides showing histological differences between sham and UPEC-infected mice at day 1, day 5 and day 10 post infection. Scale bar 50 µm. **(E and F)** Pearson correlation plot of infection time point (days post infection) and disease score of caput (E) and cauda (F). The average ± SEM disease score per time point (n=4 per time point) for sham and UPEC-infected mice is shown. Pearsońs correlation was considered to be statistically significant at p<0.05. **(G)** Volcano plot of differentially expressed genes (DEG) identified between sham and UPEC-infected mice within caput and cauda epididymides by RNASeq analysis. Numbers of DEG are indicated below the respective plot. Cut-off criteria: FDR ≤ 0.05, -1 < logFC > 1. **(H)** Top30 differentially expressed genes by comparing caput and cauda epididymides of sham and UPEC-infected mice. Cut-off criteria: FDR ≤ 0.05, -1 < logFC > 1. **(I)** Gene set enrichment analysis using DEG between caput and cauda epididymides of UPEC-infected mice. Cut-off critera: FDR < 0.2, Top up/down regulated gene sets based on gene ontology.

### Whole transcriptome and tissue analysis reveal fundamentally different immune responses in caput and cauda epididymides following infection

Initial examination pointed to different gene signatures in caput and cauda epididymides under physiological conditions, but examination under infectious conditions was not performed (Klein et al., 2019). We employed whole transcriptome analysis by RNA sequencing of total caput (including the IS), corpus and cauda 10 days p.i. to investigate the principal changes in the whole transcriptome of the different epididymal regions under pathological conditions *in vivo*. In line with the minimal histopathological alterations, almost no transcriptional differences were identified between the caput of sham and UPEC-infected mice (in total 5 DEG, cut-off: FDR ≤ 0.05, -1 < logFC > 1, Fig. 1G, Suppl. Fig. S2A, Suppl. Fig. S2B). Intriguingly, although the transcriptional profiles of the caput in sham and UPEC-infected mice were very similar (Suppl. Fig. S2A), upregulation of a few infection-related genes such as *S100a8, S100a9*, and *Slfn4* were indicative of the presence of UPEC in the infected caput (Suppl. Fig. S2B). In contrast, the cauda of sham and UPEC-infected mice showed considerable transcriptional differences (in total 5082 DEG, cut-off: FDR ≤ 0.05, -1 < logFC > 1, Fig. 1G, Suppl. Fig. S2C, Suppl. Fig. S2D). As shown by principal component analysis (PCA), the transcriptional changes in the corpus were intermediate compared with those in caput and cauda epididymides (Suppl. Fig. S2A), an observation that was reflected in a comparable magnitude of histopathological alterations (Suppl. Fig. S1E). To analyze principal differences, we focused on caput and cauda in subsequent studies as these regions displayed greater differences in gene expression levels and histopathology.

Compared to the cauda, the caput was highly enriched in transcripts encoding immunomodulatory factors, such as β-defensins, bactericidal permeability-increasing protein (BPI) and indoleamine 2,3-dioxygenase 1, with no changes in the high levels observed in sham and UPEC-infected mice (Fig. 1H). In contrast, compared to sham control mice, the cauda of UPEC-infected mice was characterized by an upregulation of numerous transcripts encoding pro-inflammatory mediators, including pro-inflammatory cytokines (e.g. *Il-1*α*, Il-6, Il-17*) and chemoattractants (e.g. *Ccl2, Ccl3, Ccl4, Cxcl2, Cxcl5*) as well as inflammasome-associated transcripts (e.g. *Nlrp3*, *Il1b*) (Fig. 1H, Suppl. Fig. S2C).

By grouping transcripts according to their gene ontology and pathway contribution, the cauda of UPEC-infected mice 10 days p.i. displayed upregulation of gene sets associated with fibrotic tissue remodeling and pro-inflammatory immune responses (e.g. positive regulation of NF kappa B – transcription factor activity, collagen fibril organization and positive regulation of the ERK1 and ERK2 cascade, Fig. 1I). Further pathway analyses revealed an upregulation of gene sets associated with B and T cell activation, indicating a transition from the innate to the adaptive immune response at this stage of infection within the cauda (Suppl. Fig. S2E). The caput epididymidis of sham and infected mice were enriched with gene sets related to sperm maturation (e.g. protein localization in cilium, cellular component assembly involved in morphogenesis and regulation of cilium beating frequency, Fig. 1I), indicative of normal epididymal function.

In line with the observed histopathological alterations and the transcriptional profile of the cauda of UPEC-infected mice, disease progression correlated positively with the appearance and degree of immune cell infiltration in this region (Fig. 2A, Suppl. Fig. S1D). Hence, we compared the dynamic of changes in immune cell populations within the caput and cauda of sham and UPEC-infected mice. Notably, we observed an increase in the total immune cell population (CD45^+^) in both the caput and cauda of sham mice that returned to normal at 10 days p.i. in caput, but remained elevated in cauda (Fig. 2B). This indicated that an immune response was elicited by surgery-associated trauma and ductal pressure due to the ligation of the vas deferens and in the absence of pathogens. We therefore extended our analysis of infiltrating immune cells to a timepoint up to three weeks p.i. to understand persistent tissue damage in the cauda described previously (Klein et al., 2020). In contrast to the caput, the cauda revealed a persistently increased percentage of CD45^+^ cells within single live cells above the levels seen in sham animals (Fig. 2B), a finding that positively correlated with the disease score (Fig. 2C).

**Figure 2:**
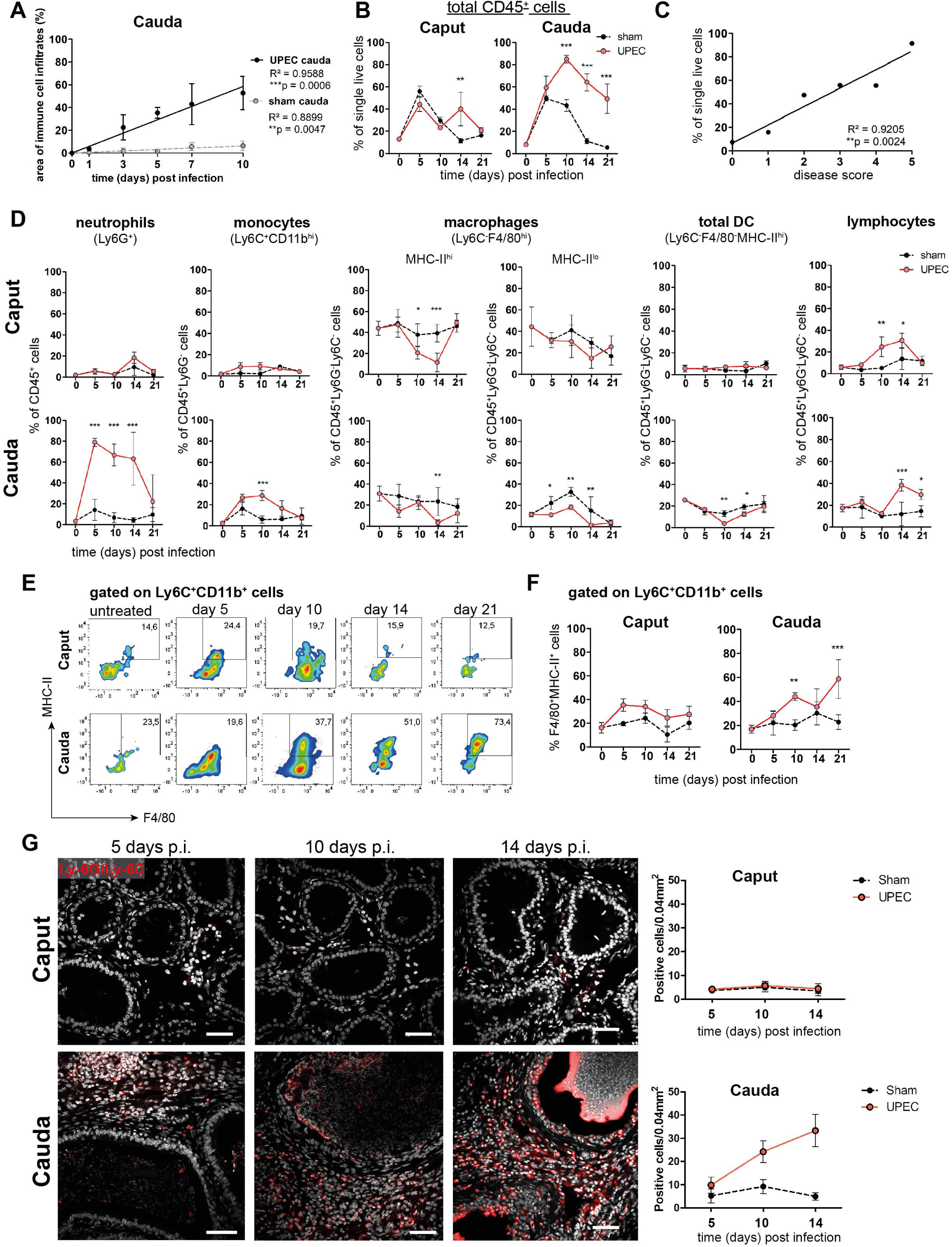
Analysis of changes in immune cell populations following infection with UPEC in C57BL/6J wild type mice. **(A)** Person correlation plot of infection time points (days post infection) and the area of immune cell infiltration within the total cauda area (%) determined by histological evaluation. Mean ± SD of at least two independent experiments with each n=4 are plotted per time point for sham and UPEC-infected mice. Pearsońs correlation was considered to be statistically significant at p<0.05 (*p<0.05, **p<0.005, ***p<0.001). **(B)** Percentage of CD45^+^ cells in single live cells within caput and cauda assessed by flow cytometry at different time points (days) post infection (mean ± SD, n=4, Two-way ANOVA with Bonferroni posthoc test, *p<0.05, **p<0.005, ***p<0.001). **(C)** Pearson correlation plot showing disease score and percentage of CD45^+^ cells in single live cells. Pearsońs correlation was considered to be statistically significant at p<0.05. **(D)** Quantification of immune cell types (as indicated above the respective diagrams) within caput and cauda at different infection time points by flow cytometry (n=4 per group per time point, mean ± SD, n=4, Two-way ANOVA with Bonferroni posthoc test, *p<0.05, **p<0.005, ***p<0.001). **(E)** Representative flow cytometry plots showing the proportion of F4/80^+^MHC-II^+^ cells within Ly6C^+^CD11b^+^ population in caput and cauda of untreated, sham and UPEC-infected mice at different time points determined by flow cytometry. **(F)** Bar diagrams showing the ratio of F4/80^+^MHC-II^+^ cells within Ly6C^+^CD11b^+^ population in caput and cauda of untreated, sham and UPEC-infected mice at different time points (mean ± SD, n=4, Two-way ANOVA with Bonferroni post hoc test, *p<0.05, **p<0.005, ***p<0.001). **(G)** Confocal microscopy images showing the location of Ly6G^+^Ly6C^+^ cells (red) within caput and cauda of UPEC-infected mice 5, 10 and 14 days post infection (nuclei in gray) including bar diagrams showing the semi-quantified summary of all immunostained tissues (by counting Ly6G^+^Ly6C^+^ cells within caput and cauda of sham and UPEC-infected mice, n=4, for each biological replicate three representative areas were counted, mean ± SD). Scale bar 50 µm.

The composition of different immune cell populations changed dynamically with disease progression (Fig. 2D, gating outlined in Suppl. Fig. S1G): in association with the initial bacterial appearance, a massive infiltration of neutrophil granulocytes (Ly6G^+^ cells) was observed, mainly in the cauda epididymidis (Fig. 2D, Suppl. Fig. S1F). Infiltration of neutrophils was followed by an influx of monocytes (Ly6C^+^CD11b^+^, Fig. 2D). Intriguingly, a significantly increased proportion of F4/80^+^CD64^+^MHC-II^+^ cells within the Ly6C^+^CD11b^+^ population was seen within the cauda from day 10 p.i. onwards, indicating that these cells were undergoing a transition into a pro-inflammatory macrophage phenotype (Fig. 2E,F). Although the overall percentage of Ly6C^+^CD11b^+^ monocytes within the CD45^+^Ly6G^-^ population returned to levels comparable to untreated mice at day 21 post infection (Fig. 2D), UPEC-infected mice showed a significantly higher ratio of F4/80^+^CD64^+^MHC-II^+^ cells within the Ly6C^+^CD11b^+^ population compared to untreated and sham control mice. As seen by immunofluorescence analysis, numbers of Ly6G^+^ and Ly6C^+^ cells were progressively increasing within the interstitium of the cauda, but not the caput, from day 10 p.i. onwards, and also became evident in the epididymal epithelium (Fig. 2G) at a time when epithelial integrity was disturbed.

Compared to sham mice, ratio of resident macrophages (Ly6C^-^F4/80^+^CD11b^+^), i.e. MHC-II^hi^ cells, tended to decrease within UPEC-infected mice during the acute phase of infection compared to sham mice in both caput and cauda (Fig. 2D). In addition, F4/80^-^MHC-II^hi^ cells, broadly considered as resident dendritic cells, also appeared to decrease within the cauda, whereas no changes were detected in the caput epididymidis (Fig. 2D). From day 10 p.i., a significant increase of cells in both caput and cauda of UPEC-infected mice was observed that were characterized by high levels of CD45 with concomitant lack of expression of Ly6G, Ly6C, CD11b, F4/80 and MHC-II - thus, these cells were broadly considered as lymphocytes without further discrimination. The increase in this population was consistent with the observed enrichment of gene sets associated with B and T cell activation at day 10 p.i. (Suppl. Fig. 2E), thus strengthening the assumption that a transition from the innate to the adaptive immune response occurred at day 10 p.i.

### Simultaneous exposure to an inflammatory stimulus *in vitro* results in differential immune responsiveness of the epididymal regions

To examine whether the observed differential immune responses within caput and cauda were merely a consequence of microbial ascension and thus the longer exposure of the cauda to the pathogens, we have utilized an *ex vivo* organ culture model that allows simultaneous challenge with an inflammatory stimulus. Cytokine production profiles of the different epididymal regions (IS, caput, corpus and cauda) were analyzed separately after stimulation with ultrapure lipopolysaccharide (LPS). While the IS and caput were still mostly unreactive, both corpus and cauda showed a significant upregulation of IL-1α, IL-1β, TNFα, MCP-1 (CCL2), IL-6 and IL-10 (Suppl. Fig. S2F). Intriguingly, IS and caput showed a higher intracellular bacteria load compared to corpus and cauda after *ex vivo* co-culture of organ pieces with UPEC, indicative for a higher and faster bacterial uptake and clearance potential (Suppl. Fig. S2G). Overall, these data suggest that the fundamentally different immunological responses observed *in vivo* within different regions of the epididymis are an inherent feature of the region, and thus independent of the administration route of the inflammatory stimulus.

### Single-cell transcriptomic analysis of immune cells in the epididymis demonstrates regional heterogeneity in steady-state

The above described observations indicated the possibility of differential immunological landscapes in the epididymal regions. To gain a comprehensive understanding, we employed single cell RNA sequencing of extravascular CD45^+^ cells (in total 12,966 cells) that were separately isolated from the four main epididymal regions (IS, caput, corpus, cauda). The data were subsequently combined into a single data set to investigate their regional distribution (Fig. 3A, Suppl. Fig. S3A-C). Unsupervised clustering and uniform manifold approximation and projection (UMAP) identified 13 different clusters (Fig. 3B) with distinct gene expression profiles (Fig. 3C). The identity of each cluster was annotated manually based on key marker gene expression (Fig. 3D, E, Suppl. Fig. S3D). Among the clusters, the majority of identified immune cells comprised several myeloid cell populations comprised several myeloid cell populations and included subsets of macrophages (clusters 1, 2 and 7), monocytes (clusters 8 and 10) and dendritic cells (clusters 2, 5 and 13). Macrophages were broadly identified by the co-expression of multiple key marker genes such as *C1qa, Fcgr1* (encoding CD64)*, Adgre1* (encoding F4/80) and *Cd68,* with alternating levels of markers such as *Cx3cr1, Ccr2, H2-Aa* (encoding a MHC-II component). Monocytes were broadly characterized by the expression of *Ly6c2, Ccr2*, and *Ace*. Dendritic cells (DC, *Flt3^+^* and high expression levels of MHC-II transcripts) were segregated into three clusters that were identified as conventional DC 1 (*Clec9a*^+^*Irf8^+^*), conventional DC 2 (*Cd209a^+^*) as well as a small population of migratory DC (*Ccr7^+^,* Fig. 3B-E). Apart from myeloid cells, all epididymal regions were populated by lymphocytes, including T cells (*Cd3e^+^*, clusters 4 and 9), NK cells (*Nkg7^+^Eomes^+^*, cluster 6) and B cells (*Cd79a*^+^, cluster 11, Fig. 3D,E). T cells were further discriminated into αβ and γδ T cells based on their alternating expression of *Trbc* and *Trdc*, respectively.

**Figure 3:**
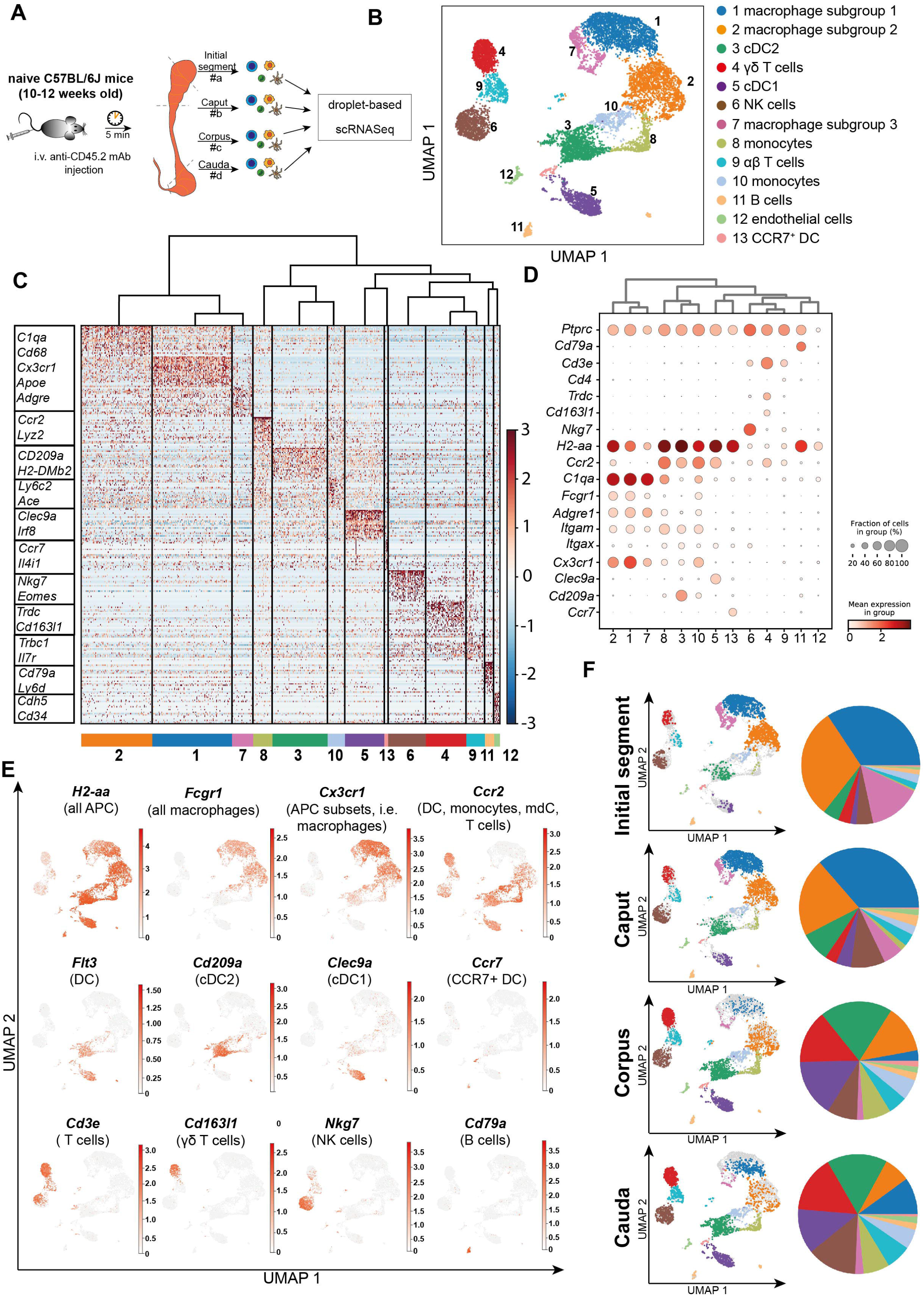
scRNASeq of different epididymal regions reveals immune cell heterogeneity within the murine epididymis under physiological conditions. **(A)** Schematic overview of the experimental procedure for isolating extravascular CD45^+^ cells from different epididymal regions. **(B)** UMAP plot of 12,966 FACS-sorted CD45^+^ cells isolated from the four epididymal regions, showing immune cell populations identified by unsupervised clustering. **(C)** Heatmap of the Top45 marker by stringent selection of markers (only present in one cluster, 585 in total) showing expression differences among clusters. **(D)** Dot plot corresponding to the UMAP plot showing the expression of selected subset-specific genes – dot size resembles the percentage of cells within the cluster expressing the respective gene and dot color reflects the average expression within the cluster. **(E)** UMAP plots showing the expression of selected key marker for the indicated immune cell population (APC-antigen-presenting cells, mdC-monocyte-derived cells, DC-dendritic cells). **(F)** UMAP plots and pie charts showing regional distribution of identified clusters.

We next defined the cluster distribution across epididymal regions (Fig. 3F, Suppl. Fig. S3D). Transcriptomic data of identified immune cell populations and their ratios within the CD45^+^ population in different epididymal regions were subsequently confirmed at the protein level by flow cytometry (Fig. 4, gating strategies in Suppl. Fig. S4). The vast majority of resident immune cells in the epididymis were found in the IS (approximately 10%-15% CD45^+^ cells among the single live cells vs. 1%-5% CD45^+^ cells in single live cells in caput to cauda (Fig. 4A). Overall, we noted similarities in the composition of resident immune cell populations in the IS and caput that were clearly distinct from that in the more distal corpus and cauda. In this regard, IS and caput were predominantly populated by macrophage subsets (approximately 78% and 66% in CD45^+^ cells, respectively (Fig. 4B) with other leukocytes accounting for < 5% for each population (Fig. 4B-H). In contrast, the corpus and cauda contained a more heterogeneous immune cell network, including several myeloid cell populations. In addition to macrophages (25%-35%, Fig, 4B), as well as monocytes (7%-10%, Fig. 4C) and dendritic cells (cDC1 7%-10%, and cDC2: 12%-20%, Fig. 4D, E) were predominantly found in close conjunction with the epididymal duct, as detected by immunofluorescence analysis. Lymphocyte subsets (NK cells [10%, Fig 4F], B cells [2-5%, Fig. 4G], T cells [10-20%, Fig 4H]) were located in both the interstitial and intraepithelial compartment (Fig. 4F-H). Among the T cells, we further distinguished αβ and γδ T cells, with only the latter found within the epithelium (Fig. 4H). The difference in leukocyte populations, their ratio and tissue localization throughout the epididymis points to the existence of inherently different immunological environments in the proximal (IS, caput) and distal regions (corpus, cauda), which form the basis of the differential immune responsiveness observed in models of epididymitis.

**Figure 4:**
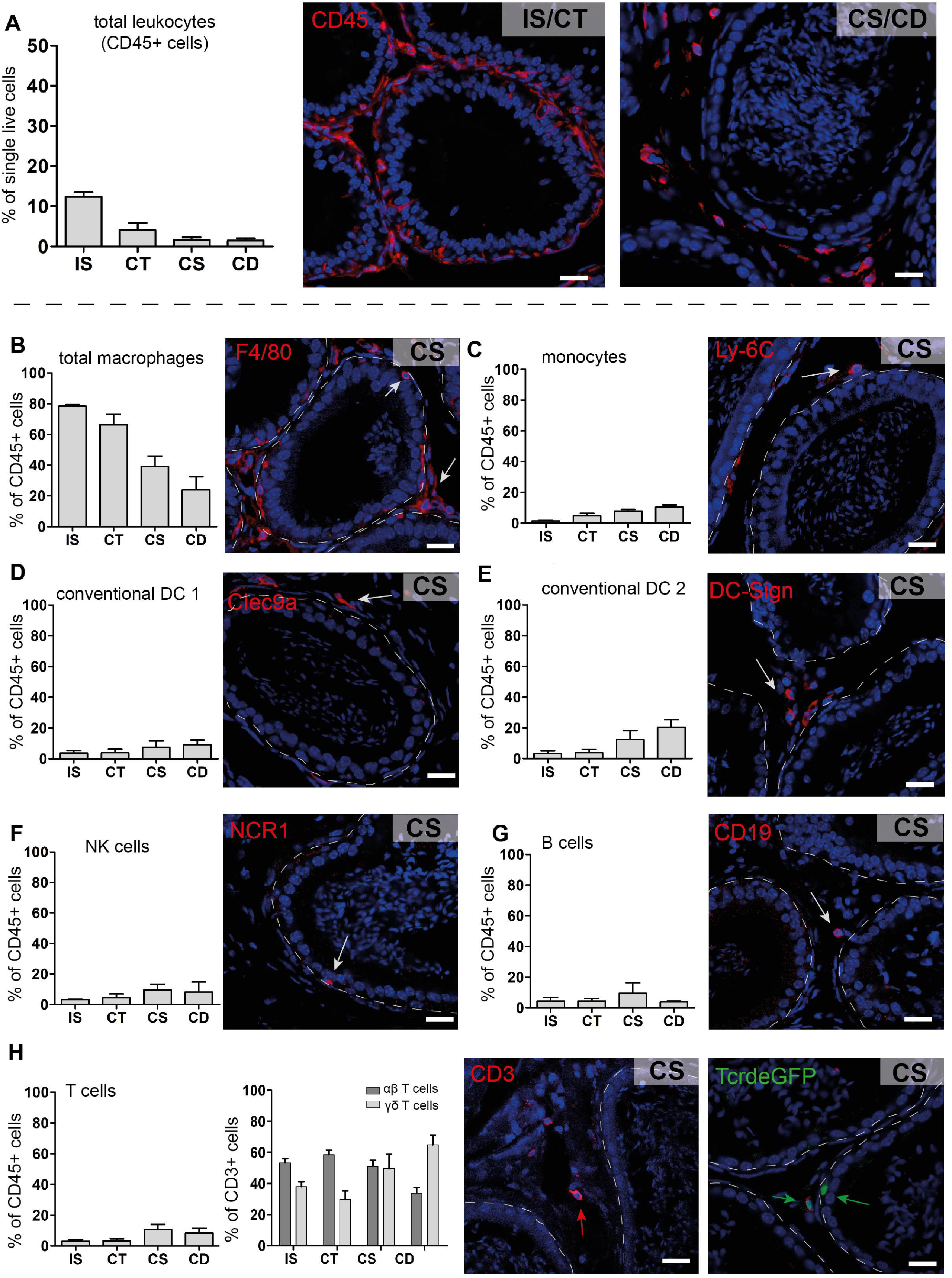
Quantification and localization of identified immune cell populations among epididymal regions (Scale bar 20 µm). **(A)** Distribution (assessed by flow cytometry n=4-8, bar diagram showing mean ± SD) of total leukocytes (CD45^+^ cells) and localization within proximal (IS/CT) and distal (CS/CD) regions, shown by immunostaining of CD45. **(B-H)** Quantification and localization of the following immune cell populations were assessed by flow cytometry and immunostaining using selected marker (n=4-8, mean ± SD). Representative images are displayed from the corpus (CS) regions: **(B)** Total macrophages (F4/80^+^, red), located in the interstitial, intraepithelial and peritubular compartments, **(C)** Monocytes (Ly-6C^+^), located in the peritubular compartment, **(D and E)** Conventional dendritic cells cDC 1 (Clec9a^+^) and 2 (DC-Sign/CD209a^+^), **(F)** NK cells (NK1.1^+^ for flow cytometry and NCR1 for immunostaining), located in the intraepithelial compartment, **(G)** B cells (B220/CD45R^+^ for flow cytometry and CD19^+^ for immunostaining), **(H)** T cells that were further segregated into αβ T cells (TCRβ^+^, red) and γδ T cells (TCRγδ^+^, green).

### Macrophages separate into several subgroups based on their transcriptional profile

*Adgre1^+^C1qa^+^* cells, broadly considered as epididymal macrophages, constitute the majority of CD45^+^ cells in the epididymis. In subsequent closer analyses with the aim to decipher the possible heterogeneity of this population, we firstly distinguished macrophage populations (clusters 1, 2 and 7) from monocyte populations (clusters 8 and 10) based on their expression of *C1qa*, *Ccr2*, *Ly6c2*, *Napsa* and *Plac8* (Fig. 5A), with both monocyte clusters expressing lower levels of *C1qa.* However, cluster 10 showed higher expression of transcripts encoding classical monocyte markers (*Ly6c2, Napsa, Plac8*) compared to cluster 8. This indicated that cluster 10 resembles a classical monocyte population, whereas cluster 8 represents a monocyte population undergoing differentiation into a macrophage phenotype. This assumption was also supported by intermediate expression of *C1qa, Adgre1* and *Fcgr1* between classical monocytes (cluster 10) and macrophage populations (cluster 1, 2 and 7, Fig. 3D, Fig. 5A).

**Figure 5:**
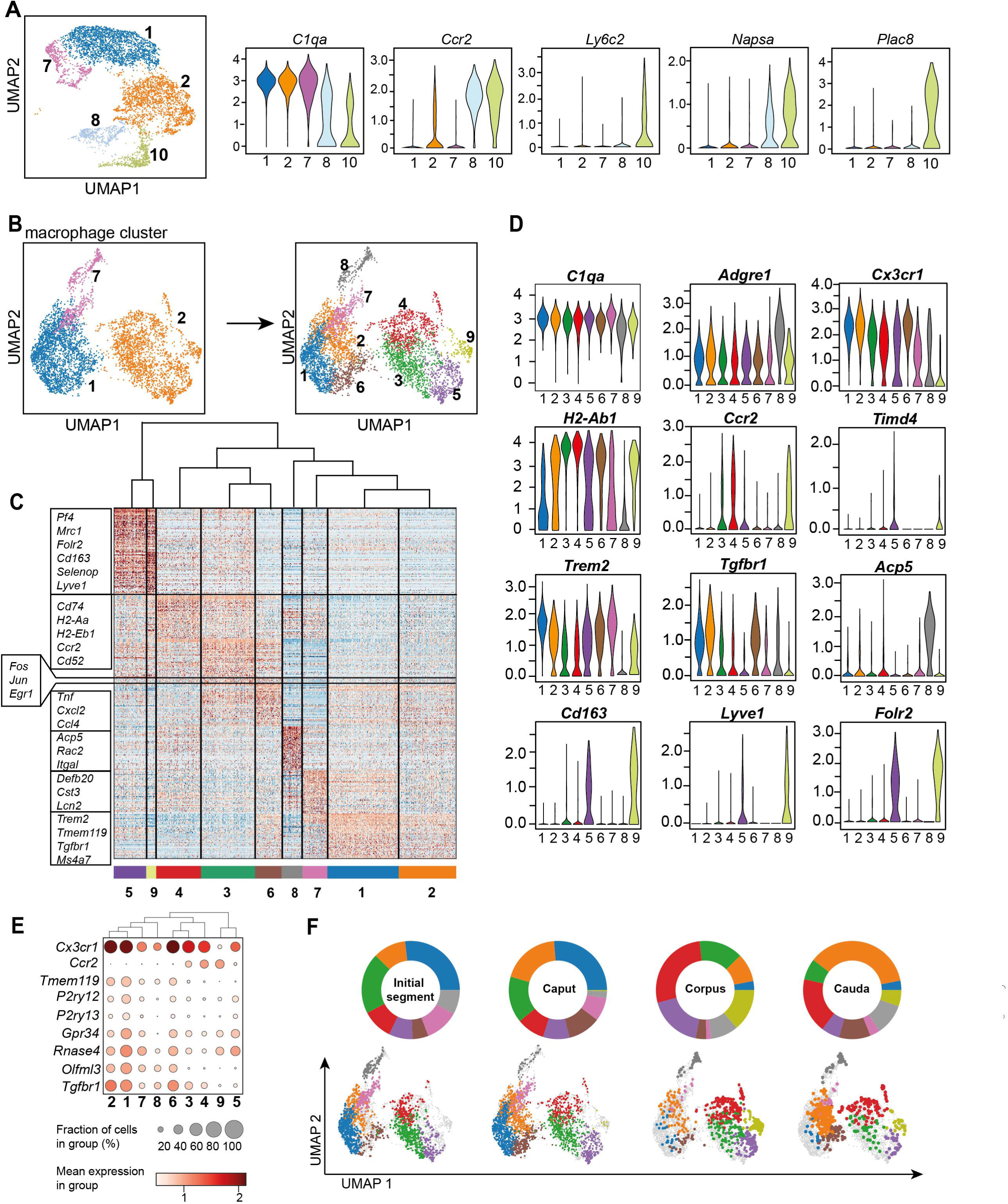
Distinct macrophage subgroups exist within the murine epididymis. **(A)** UMAP plot and violin plots showing segregation of macrophages (clusters 1, 2, 7) and monocytes (clusters 8, 10) based on clustering and expression of the selected key genes *C1qa, Ccr2, Ly6c2, Napsa, Plac8*. **(B)** UMAP plot showing re-clustering of macrophage population (clusters 1, 2, 7) under exclusion of all other previously identified CD45^+^ cluster resulting in the formation of nine *Adgre1^+^* subclusters. **(C)** Heatmap of the 50 most differentially expressed marker genes in each cluster from Fig. 4B. **(D)** Violin plots showing the expression level of selected genes. **(E)** Dot plot corresponding to the UMAP plot showing the expression of selected subset-specific genes – dot size resembles the percentage of cells within the cluster expressing the respective gene and dot color reflects the average expression within the cluster. **(F)** UMAP plots and pie charts showing the distribution of identified macrophage populations among epididymal regions.

To further characterize the heterogeneity among macrophage subpopulations (clusters 1, 2 and 7), *Adgre1*^+^*C1qa^+^* cells were re-analyzed after exclusion of other CD45^+^ cells. By unsupervised clustering, nine subgroups (Fig. 5B) were identified, each with a distinct gene expression profile (Fig. 5C, Suppl. Fig. S5A). All identified macrophage subgroups were highly enriched with *C1qa* and *Adgre1* transcripts confirming their macrophage identity (Fig 5D, Fig. S5A). Clusters 1 and 2 constitutes the majority of macrophages and demonstrated comparatively high expression levels of genes that were previously reported to be associated with homeostatic and sensing functions of macrophages within other tissues (i.e. brain microglia, (Hickman et al., 2013; van Hove et al., 2019; Abels et al., 2021), indicating similar functions for the epididymis. These genes include *Cx3cr1, Tmem119, P2ry12, P2ry13, Gpr34, Rnase4, Olfml3* and *Tgfbr1* (Fig. 5D, Fig. 5E). In contrast to cluster 1, cluster 2 expressed high levels of MHC-II component transcripts (e.g. *H2-Ab1*, Fig. 5D), indicating an activated status for antigen-presentation. Cluster 6 showed a similar expression pattern to cluster 2, but was highly enriched with transcripts encoding several cytokines and chemokines (*Tnf, Cxcl2, Ccl4*), as well as immediate-early response genes, such as *Fos, Jun* and *Egr1* (Fig. 5C and Suppl. Fig. S5B). However, it is likely that the latter genes could be a consequence of tissue processing prior to sequencing (Denisenko et al., 2020). These transcriptional differences indicate that cluster 6 constitutes an activated form of cluster 2, hence, both clusters were considered as one subgroup.

Clusters 3 and 4 were enriched with *Ccr2* and transcripts encoding MHC-II components (e.g. *H2-Ab1, H2-Aa, H2-Eb1,* Fig. 5D), indicating an activated pro-inflammatory phenotype. Albeit transcriptionally similar to cluster 4, cluster 3 expressed high levels of several activation genes, such as immediate-early response genes (*Fos, Jun, Egr1*, Fig. 5C, Suppl. Fig. S5B) and cytokine and chemokines (Fig. 5C), suggesting that cluster 3 also constituted an activated subset, as was the case for cluster 6, consequently, cluster 3 and 4 were also considered as one subgroup. Cluster 5 and 9 were both enriched with *Cd163, Lyve1* and *Folr2*, but showed reciprocal expression of *Timd4* and *Ccr2*, respectively (Fig. 5D, Suppl. Fig. S5A), indicating similar anti-inflammatory or regulatory phenotypes, but different ontogenies. Clusters 7 and 8 constituted rather minor subgroups and did not express either *Ccr2* or *Cd163, Lyve1* or *Timd4,* and only low levels of *Cx3cr1* compared to the other subgroups. In contrast to Cluster 8, Cluster 7 cells expressed *Trem2* beside MHC-II encoding transcripts such as *H2-Ab1* (Fig. 5D). Cluster 8 showed a relatively high expression level of *Adgre1* compared to all other clusters in addition to high levels of *Acp5* (Fig. 5D). The differential distribution among regions was the most striking difference among the identified subpopulations Fig. 5F).

### Macrophage subgroups show striking regional differences in their regional and compartmental distribution

Based on the transcriptional profiles, seven macrophage subpopulations (subgroup 1, [2+6], [3+4], 5, 7, 8, 9) were distinguished in the murine epididymis. In the next step, identified subpopulations were quantified in support by flow cytometry in wild type mice and localized in the tissue using immunofluorescence in *Cx3cr1*^GFP^*Ccr2*^RFP^ reporter mice. Overall, F4/80^+^ cells constituted approximately 80% of CD45^+^ cells within the IS and these cells gradually decreased towards the cauda to approximately 25% of CD45^+^ cells (Fig. 6A). F4/80^+^ cells were found throughout all epididymal regions to be constituents in both epididymal compartments, i.e., the ductal epithelium and the interstitium. The majority of F4/80^+^ cells were also CX3CR1 positive (Fig. 6B and 6C, Suppl. Fig. S6A). Only a small fraction of intraepithelial CX3CR1^+^ cells was F4/80^-^ within the IS (indicated by arrowheads Fig. 6C, Suppl. Fig. S6A).

**Figure 6:**
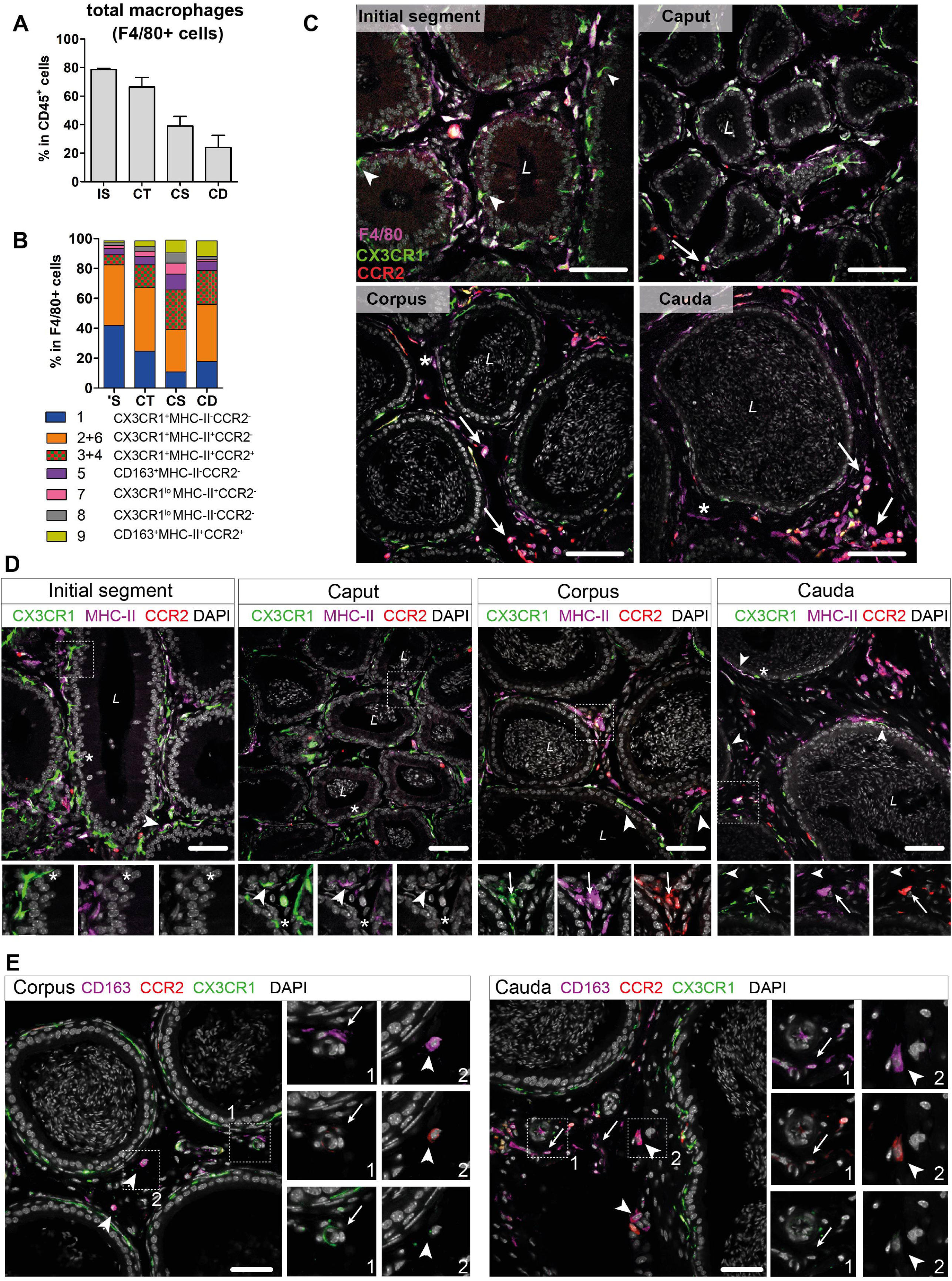
Distribution and localization of identified macrophage subgroups by flow cytometry and immunofluorescence. **(A)** Bar diagram showing the percentage of F4/80^+^ cells within the CD45^+^ population throughout the epididymal regions, assessed by flow cytometry (n=8, mean ± SD). **(B)** Stacked bar diagram displaying the percentages of identified macrophage subtypes within the F4/80^+^ population throughout epididymal regions assessed by flow cytometry. Markers were selected based on scRNASeq results (n=4). **(C)** Confocal microscopy images of F4/80 staining (purple) on *Cx3cr1*^GFP^*Ccr2*^RFP^ adult reporter mice. The majority of interstitial and intraepithelial CX3CR1^+^ cells were F4/80^+^. Arrowheads indicate a small fraction of intraepithelial CX3CR1^+^ F4/80^-^ cells found within the IS. Arrows indicate interstitial F4/80^+^ cells that were CX3CR1^-^ and CCR2^+^ within caput, corpus and cauda epididymides. Asterisks (*) label a small fraction of F4/80 single positive cells (CX3CR1^-^CCR2^-^) found in the corpus and cauda. Scale bar 50 µm (L = Lumen). **(D)** Confocal microscopy images of MHC-II staining (purple) on *Cx3cr1*^GFP^*Ccr2*^RFP^ adult reporter mice. Asterisks (*) indicate intraepithelial CX3CR1^+^MHC-II^-^ cells within the IS and caput epididymides. Arrowheads indicate CX3CR1^+^MHC-II^+^ cells, lining the epididymal duct within the IS and situated within the epithelium within caput, corpus and cauda epididymides. Arrows indicate interstitial CX3CR1^+^MHC-II^+^CCR2^+^ cells additionally found within corpus and cauda epididymides. Scale bar 50 µm (L = Lumen). **(E)** Confocal microscopy images of CD163 staining (purple) on *Cx3cr1*^GFP^*Ccr2*^RFP^ adult reporter mice in corpus and cauda epididymides. Arrows indicate CD163 single positive cells that were found in close proximity to vessels within the corpus and cauda. Arrowheads indicate CD163^+^CCR2^+^ cells found solitarily distributed within the interstitium in the corpus and cauda. Scale bar 50 µm.

In accordance with the scRNASeq data, CX3CR1^+^CCR2^-^MHC-II^-^ macrophages (cluster 1) and CX3CR1^+^CCR2^-^MHC-II^+^ (clusters 2 and 6) macrophages constituted the majority of F4/80^+^ cells and both were highly abundant within the IS (40% of total F4/80^+^ cells for each population, Fig. 6B). Both subgroups declined towards the cauda (Suppl. Fig. S5C), although the ratio of CX3CR1^+^CCR2^-^MHC-II^+^ cells (cluster 2) within the F4/80^+^ population remained similar throughout all epididymal regions (Fig. 6B). The majority of CX3CR1^+^CCR2^-^MHC-II^-^ macrophages (cluster 1) were located within the epididymal epithelium (Fig. 6C, Fig. 6D), between adjacent principal cells. Notably, while intraepithelial macrophages within the IS, which exhibited long and thin protrusions towards the lumen, did not express MHC-II, these intraepithelial cells gained MHC-II expression in caput, corpus and cauda (indicated by arrowheads in Fig. 6D and Suppl. Fig. S6B). In addition, CX3CR1^+^CCR2^-^MHC-II^+^ cells (clusters 2 and 6) closely surrounded the epididymal duct with highest density in the IS (Fig. 6D and Suppl. Fig. S6B).

In contrast, CX3CR1^+^CCR2^+^MHC-II^+^ macrophages (clusters 3 and 4) showed the opposite distribution pattern. While being less abundant in the IS and caput (5%-10%), CX3CR1^+^CCR2^+^MHC-II^+^ macrophages constituted 20%-30% of macrophages in the corpus and cauda (Fig. 6B), a similar proportion to the CX3CR1^+^CCR2^-^MHC-II^+^ macrophages (cluster 2). Notably, triple positive CX3CR1^+^CCR2^+^MHC-II^+^ macrophages (clusters 3 and 4) were localized exclusively in the interstitium, most prominently in the cauda (Fig. 6D, Suppl. Fig. S6B).

Apart from these three major populations, the minor populations were further subdivided based on the expression of CD163 and CCR2 (Suppl. Fig. S4D). Here, CD163^+^CCR2^-^ macrophages (cluster 5) constituted 3%-10% of resident macrophages and were predominantly found in the corpus (Fig. 6B, Suppl. Fig. S5C). Similarly, CD163^+^CCR2^+^ macrophages (Cluster 9) were most abundant in the corpus and cauda (10% of total F4/80^+^ cells, Fig. 6B, Suppl. Fig. S5C). In tissue sections, both populations were located interstitially, whereby CD163 single positive cells (CCR2^-^) were clustered in close proximity to vascular structures and appeared smaller in size compared to the solitarily distributed CD163^+^CCR2^+^ cells (Fig. 6E). Cells that were F4/80^+^ but CX3CR1^-^ concomitant with the absence of CCR2 and CD163 were considered to be macrophages of cluster 7 and 8. Furthermore, cells belonging to cluster 7, but not cluster 8, express MHC-II. Both populations were most abundant within the corpus (Fig. 6B), but constituted only a small fraction of resident immune cells (approximately 1%-4% of total CD45^+^ cells, Suppl. Fig. S5C). Generally, F4/80^+^ cells negative for both CX3CR1 and CCR2 (belonging to both cluster 7 and 8) were exclusively found within the interstitium (indicated by an asterisk in Fig. 6D and Suppl. Fig. S6B).

Overall, the regional and compartmental distribution suggests that distinct macrophage subsets populate the epididymis to facilitate the complex spectrum of canonical macrophage functions (homeostatic, inflammatory, reparative/ regulatory) adapted to the needs of the respective microenvironment: ‘Scavenger functions’ within the epithelial compartment of the proximal regions (i.e. IS) to maintain tissue integrity, and ‘guardian functions’ within the distal regions to efficiently tackle invading pathogens and tissue regeneration.

### Maintenance of resident macrophages in epididymal regions depends differentially on monocyte recruitment

Having identified the variation and heterogeneity of resident macrophages among epididymal regions, we further investigated putative differences in the monocyte-dependence on the maintenance of resident macrophages among the epididymal regions. Parabiosis experiments were performed by surgically conjoining CD45.1^+^ wild type mice with CD45.2^+^ *Ccr2^-/-^* mice for 6 months before analyzing the ratio of monocyte-derived CD45.1^+^ cells within Ly6C^hi^ blood monocytes (Fig. 7A. Suppl. Fig. S7A) and resident macrophage populations of the *Ccr2^-/-^* recipient mouse (Fig. 7A). Within the blood, approximately 40%-60% of Ly6C^hi^ cells originated from the CD45.1 donor, indicating efficiently established chimerism. For the epididymis, all CD11b^+^CD64^+^ macrophages (containing all previously identified subgroups) were gated and further subdivided using CCR2 and TIMD4, respectively (Fig. 7B, Suppl. Fig. S7B), as markers for monocyte-derived and self-maintaining macrophages, as previously described in other organs (Dick et al., 2022). Epididymal fat was additionally investigated to examine if a directly associated neighboring tissue differs with respect to the monocyte contribution. All CCR2^+^ cells (corresponding to cluster 3, 4 and 9), which are most abundant within the corpus and cauda (approximately 30-40 cells/ mg tissue, Fig. 7D), were exclusively of donor origin in all regions (Fig. 7C and 7G). In contrast, TIMD4^+^ cells (corresponding to cluster 5) that were found in low numbers in corpus and cauda (approximately 40-50 cells per mg tissue, Fig.7E) were CD45.1^-^ indicating that this population did not originate from the donor (Fig. 7C and 7H). The majority of CD64^+^CD11b^+^ cells were CCR2^-^TIMD4^-^ (double negative) and were most abundant in the IS (Fig. 7F). This population displayed the majority of resident macrophages within the epididymis (all previously described subpopulations, mainly CX3CR1^hi^ macrophages cluster 1 and 2, but also CX3CR1^lo^ subpopulations clusters 7 and 8). Intriguingly, although double negative epididymal macrophages were generally less monocyte-dependent compared to macrophages located within the epididymal fat (Fig. 7C and 7I), significant differences were detected between the proximal (IS, caput) and distal regions (corpus, cauda) of the epididymis. While CCR2^-^TIMD4^-^ macrophages within the IS and caput had only very low donor chimerism (approximately 5%-10% [normalized to blood, comparable to TIMD4^+^ macrophages), macrophages from corpus and cauda showed a much higher chimerism (30%-40% [normalized to blood], Fig. 7C and 7I).

**Figure 7:**
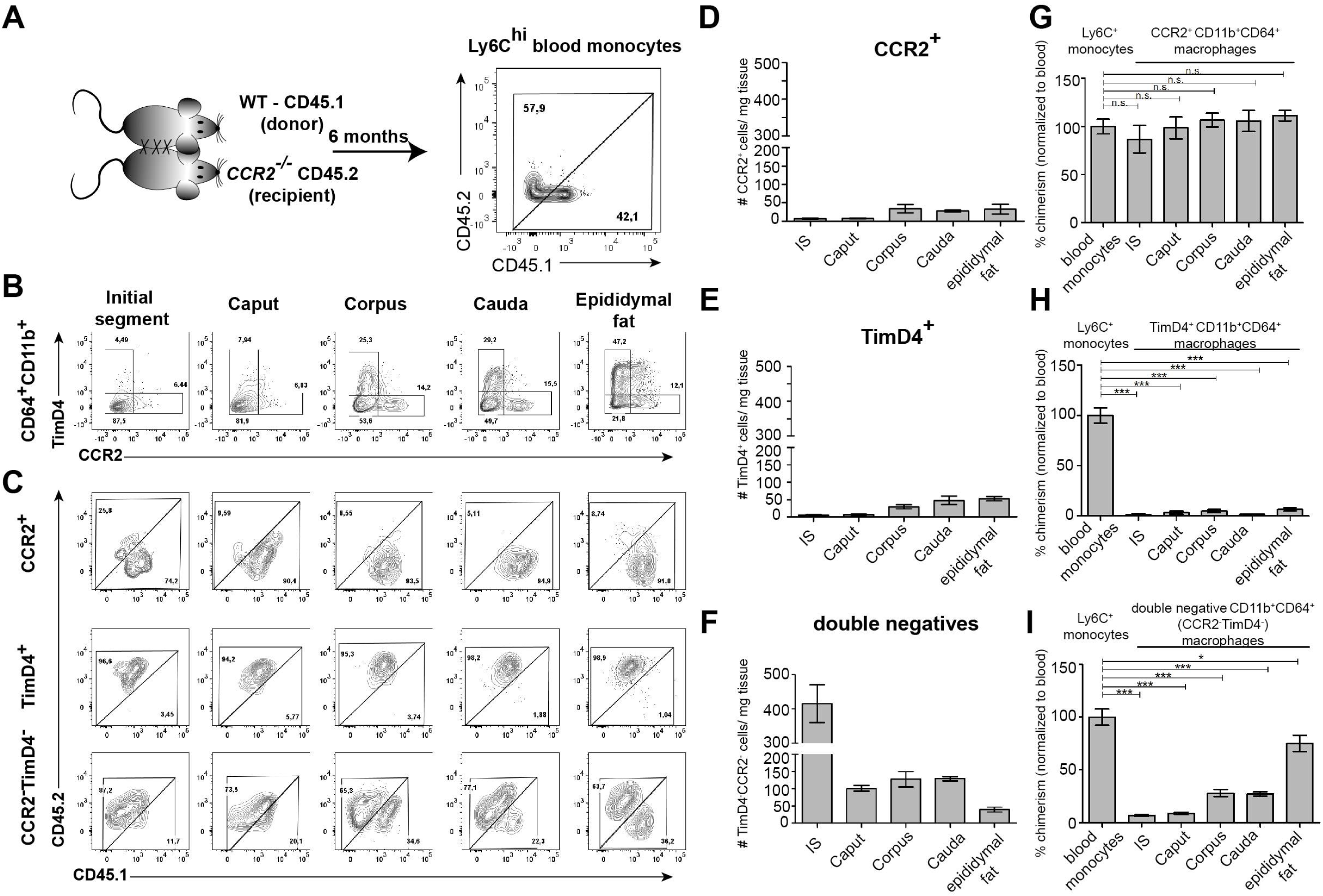
Resident macrophages differentially depend on monocytes within epididymal regions. **(A)** Parabiosis was conducted by surgically conjoining wild type CD45.1^+^ donor mice with CD45.2 recipient *Ccr2^-/-^* mice for 6 months. Donor chimerism was confirmed on CD115^+^CD11b^+^Ly6C^hi^ monocytes. **(B)** Flow cytometry contour plots showing the segregation of resident macrophages (CD11b^+^CD64^+^) isolated from different epididymal regions using the ontogeny marker TIMD4 and CCR2. Epididymal fat served as control tissue. Plots are representative for six parabionts. **(C)** Flow cytometry contour plots showing the chimerism in CCR2^+^, TIMD4^+^ and CCR2^-^TimD4^-^ macrophages within different epididymal regions based on the CD45.1 and CD45.2 labelling. Plots are representative for six parabionts. **(D-F)** Bar diagrams showing the number of CCR2^+^ **(D)**, TIMD4^+^ **(E)** and CCR2^-^ TimD4^-^ **(F)** macrophages (CD64^+^CD11b^+^) within different epididymal regions of the analyzed recipient *Ccr2^-/-^*mice (mean ± SEM, n=6). **(G-I)** Bar diagrams showing the percentage of chimerism normalized to blood chimerism in CCR2^+^ **(G)**, TimD4^+^ **(H)** and CCR2^-^TIMD4^-^ **(I)** epididymal macrophages (CD64^+^CD11b^+^) in the recipient *Ccr2^-/-^* mice after six months (n=6, n.s. = not significant, *p<0.05, **p<.0.005, ***p<0.001, n=6, mean ± SEM, one-way ANOVA with Bonferroni multiple comparison test)

In regional terms, these data indicate a higher monocyte-dependent turnover rate of resident macrophages within the distal epididymis (corpus and cauda) in which ascending pathogens enter the epididymis first, but proposes only a minor impact of monocytes in the maintenance of macrophages within the proximal regions (IS and caput).

## Discussion

In spite of the fact that previous studies clearly showed that the different epididymal regions develop distinct immune responses following bacterial infection (Michel et al., 2016; Klein et al., 2020), the underlying mechanisms remained elusive. In our initial experiments, we confirmed findings from previous studies showing that immunopathological damage following UPEC infection occurs almost exclusively in the cauda epididymidis (Michel et al., 2016; Klein et al., 2020) with loss of epithelial integrity, interstitial fibrosis and duct obstruction. Expanding on these previous observations, we further demonstrate that the accompanying leukocytic infiltration is characterized by a massive influx of neutrophils and monocyte-derived MHC-II^hi^ macrophages with concurrent loss of epithelial integrity. Both neutrophils and monocyte-derived macrophages are important first defenders of the innate immune response during acute infection due to their high phagocytic activity. However, during immune response against microbes both populations can also elicit substantial collateral tissue injury by releasing inflammatory and cytotoxic mediators that, in turn, amplify the immune response (Segel et al., 2011; Kruger et al., 2015). The role of neutrophils and monocyte-derived macrophages in tissue injury in the cauda has become evident in mice lacking CCR2, which is required for recruitment of circulating immune cells to inflammatory sites. *Ccr2^-/-^* mice show a significantly reduced influx of neutrophils and inflammatory monocyte-derived macrophages concomitant with less severe tissue damage in the cauda during UPEC infection compared to wild type mice (Wang et al., 2021) pointing to a role as double-edged swords in acute epididymitis by participating both in defense and inflammation-associated tissue damage. Nevertheless, bacterial virulence factors may contribute to some extend to tissue damage in the cauda following the observation that UPEC persist in this region much longer and at higher numbers than in the other parts of the organ. A further driving force of the persisting immunopathology seen in the cauda could relate to the extravasation of immunogenic spermatozoa through the damaged epithelial barrier, which may trigger accumulated MHC-II^hi^ macrophages and lymphocytes (B and T cells) towards an adaptive immune response against spermatozoal neo-antigens. This is supported by pathway analysis of gene sets associated with B and T cell activation and granuloma formation indicating a transition from innate to adaptive immune response limited to the cauda. Of note, granulomas can be induced by interstitial sperm injection alone also leading to massive tissue destruction in the cauda epididymidis (Itoh et al., 1999) and formation of anti-sperm antibodies as seen in another model of *E. coli*-elicited epididymitis and in epididymitis patients (Ingerslev et al., 1986; Nashan et al., 1993; Lotti et al., 2018; Silva et al., 2021).

Contrasting to the strong pro-inflammatory processes within the cauda, the caput remains mostly unaffected – an observation that previously raised the question whether the caput is either non-responsive or to a lesser extent responsive compared to the cauda. As bacteria are initially present in the caput, albeit at lower numbers and for a shorter time (potentially due to a faster bacterial clearance potential as evindenced by the *ex vivo* approach), it can be excluded that the lower bacterial load is an explanation for the differential immune response in the caput. The very mild and transient immune response in the caput is characterized by the upregulation of a very small number of genes that are indicative of a limited inflammatory response triggered by the pathogens, such as the alarmins *S100a8* and *S100a9.* Interestingly, both alarmins have previously been demonstrated to be upregulated within the kidney and bladder during UPEC-elicited urinary tract infection, but did not substantially contribute to an effective host immune response (Dessing et al., 2010). Possibly, the upregulation of *S100a9* within the caput could drive macrophages to polarize to an anti-inflammatory and immunosuppressive phenotype as seen in the testis following UPEC infection (Fan et al., 2021). A clear indication for a regionalized immune response with a predominant reaction in the cauda is derived from this and other studies that use an inflammatory stimuli such as LPS that act simultaneously on all regions *in vivo* and *in vitro* rather than gradually ascending such as an *in vivo* UPEC infection (Supplemental Figure S2F, (Silva et al., 2018; Wang et al., 2019; Wijayarathna et al., 2020).

Taken together, it is evident that fundamental differences must exist in the immunological milieus of the epididymal regions that most likely rely on leukocyte subpopulations that gradually change in phenotype throughout the organ. Initial evidence came from previous studies (Da Silva et al., 2011; Da Silva and Smith, 2015; Da Silva and Barton, 2016; Voisin et al., 2018; Battistone et al., 2020). However, the full extent of the heterogeneity of resident immune cells and their identity remained elusive. Intriguingly, our data reveal that distinct immunological landscapes exist within proximal (IS, caput) and distal regions (corpus, cauda), that are tailored to the respective needs of the microenvironments.

Overall, macrophages constitute the major immune cell population of the murine epididymis and exhibit a dense network consisting of several transcriptional distinct subpopulations that populate different niches according to their homeostatic, reparative and inflammatory properties. CX3CR1^hi^ macrophages that possess a homeostatic and sensing profile are situated within and around the epididymal epithelium with highest abundance in the IS. These macrophages extend long dendrites towards the lumen, consistent with previous observations (Da Silva et al., 2011; Battistone et al., 2020). The transcriptional profile of intraepithelial CX3CR1^+^ cells combined with the known high phagocytic potential towards apoptotic epithelial cells (i.e. within the IS, (Smith et al., 2014)) and pathogens (Battistone et al., 2020) indicates a central function in tissue homeostasis and immune surveillance in order to efficiently maintain epithelial integrity. Of note, the high density of sensing CX3CR1^hi^ macrophages in combination with the narrow lumen of the IS indicates a potential function of this region as ‘immunological bottleneck’ in which the luminal content is constantly monitored in order to induce tolerance towards immunogenic sperm antigens and to eliminate pathogens from further ascend to the testis. The presence of CX3CR1^+^ cells within the IS was previously described, however, were initially related to dendritic cells (Da Silva et al., 2011) and subsequently generally as MP based on morphology and partial CD11c expression (Da Silva and Smith, 2015; Da Silva and Barton, 2016). In our study the transcriptional profile clearly indicates a macrophage phenotype.

In contrast, the distal regions (corpus, cauda) are populated by a more heterogeneous macrophage pool consisting of less intraepithelial CX3CR1^+^ macrophages, but higher abundance of interstitial pro-inflammatory monocyte-derived CCR2^+^MHC-II^+^, vasculature-associated TLF^+^ macrophages (expressing a combination of *Timd4, Lyve1, Folr2*; marker used in the present study CD163) as well as CX3CR1^-^TLF^-^CCR2^-^ macrophages (contained in clusters 7 and 8). The co-existence of these three populations was recently reported to be conserved across organs (Dick et al., 2022) and, similar to other organs, the different macrophage pools have distinct monocyte contributions. While TLF^+^ macrophages are rather self-renewing and CCR2^+^ macrophages monocyte-dependent, TLF^-^CCR2^-^ macrophages (including CX3CR1^+^ macrophages in our parabiosis experiment with alternating MHC-II levels) are partially dependent on monocytes in distal, but not in proximal epididymal regions. These findings support the conclusion that local environmental factors could influence parameters that regulate monocyte entry and replacement of distinct macrophage populations in different regions of the epididymis.

The co-existence of antigen-presenting myeloid cell populations (macrophages, monocytes, dendritic cells) with lymphocyte subtypes (NK cells, αβ T cells, γδ T cells, B cells) within the distal regions of the epididymis implies an environment of immune-responsiveness. As we found a higher proportion of monocytes-derived cells and conventional DC 1 and 2 (including a small fraction of activated *Ccr7*^+^ DC) within the distal regions, an ongoing antigen-sampling and interaction with the draining lymph node can be assumed but would require further confirmation before a better understanding of the region-specific role of migratory myeloid cells in the epididymal immune regulation can be achieved. The existence of intraepithelial and interstitial lymphocyte subpopulations with innate-like characteristics (i.e. NK cells, γδ T cells), predominantly within the distal regions, implies a contribution of these cells to the onset of immune responses against pathogens. Both populations generally function as key responders to barrier stress signals in mucosal tissues and accelerate proinflammatory processes by secreting effector cytokines, particularly IL-17- and IFNγ (Shi et al., 2011; Papotto et al., 2017). Since the function of innate lymphoid cells highly depends on their activation status and functional polarization within the periphery (Bonneville et al., 2010; Klose and Artis, 2020), local environmental factors may determine regulatory versus cytotoxic functions of NK and γδ T cells in different epididymal regions.

As a limitation, our scRNASeq approach supplies ‘only’ a snapshot of extravascular immune cells within the epididymis at a defined time (in the adult) neglecting times of residency for e.g. myeloid and lymphoid immune cell populations that possess migratory capabilities and patrol between non-lymphoid tissues and draining lymph nodes, a process required for immune surveillance and induction of immune responses or tolerance (Hampton and Chtanova, 2019).

Together, our data provide the first atlas of extravascular CD45+ cells within the murine epididymis under normal conditions. Strategical positioning of identified immune cell populations strongly indicates the existence of distinct immunological landscapes at the opposing ends of the epididymal duct that, in turn, is considered as a main driver for the observed differences in the intensity of immune responsiveness upon bacterial infection. We believe that the data in the present study provide a valuable starting point and common research platform for future studies on the organ-specific function of these populations in epididymal immunity.

## Acknowledgments

The study was supported by grants from the Deutsche Forschungsgemeinschaft (DFG), Monash University and the Medical Faculty of Justus-Liebig University to the International Research Training Group on ‘Molecular pathogenesis of male reproductive disorders’ (GRK 1871). We would like to thank Martina Hudel for preparing the UPEC suspensions and Suada Fröhlich for assistance in tissue processing.

## Author contributions

CP, DA, MH, LW, DB, DK, CK, SB carried out experiments and analyzed data (CP and SB conducted *in vivo* infection experiments (CP and DA performed flow cytometry of immune cells under physiological and pathological conditions and analyzed data; MH, DA and CP analyzed UPEC-infected mice [histology, morphometrics, disease scoring, immunostaining]; DB performed *ex vivo* organ culture experiments; LW and DK carried out localization studies). CK, SE, CP, SB performed parabiosis experiments and cell preparation for scRNASeq analyses. SG and CP performed whole transcriptome analyses of UPEC-infected tissue and data analyses of scRNASeq data. SR and JM were involved in the data interpretation about particular immune cell populations (γδ T cells, and DC, respectively) and supplied critical reagents and cellular analysis tools. CP, MPH, KLL, RM and AM conceptualized the study. CP, SE, SB, MPH, JM, SR, KLL, RM and AM were involved in writing and critical revision of manuscript. All authors gave final approval of the submitted and published version.

This study has not been published elsewhere before.

## Methods

### Mice

All mice used in this study were purchased from Charles River and Jackson Laboratories and/ or bred under pathogen-free conditions prior to use at the animal facilities of Justus Liebig-University Giessen, Germany (C57BL/6J WT (Charles River), *Cx3Cr1*^GFP^*Ccr2*^RFP^ (JAX ID: 032127, Jackson Laboratories), The Toronto General Research Institute, Canada (C57BL/6J CD45.1 (JAX ID: 002014), *Ccr2*^-/-^ (JAX ID: 004999)) and the Central Animal Facility at Hannover Medical School (Tcrd-H2BeGFP, JAX ID: 016941).

All animal experiments were approved by the respective local Animal Ethic Committees (Germany: Regierungspräsidium Giessen GI20/25 G60/2017, GI20/25 G71/2019 and Canada AUP: 4054.37). Killing of wild type C57BL/6J and *Cx3Cr1*^GFP^*Ccr2*^RFP^ mice without any prior treatment had been declared to the Animal Welfare Officer of Justus-Liebig-University Giessen, Germany (Registration No. M_684 and M_ 755, respectively). Experiments were conducted in strict accordance with the Guidelines of the Care and Use of Laboratory Animals of the German law for animal welfare, the European legislation for the protection of animals for scientific purposes (2010/63/EU) and the Guidelines of the Canadian Council of Animal Care. For euthanasia prior to organ collection, mice were deeply anesthetized by inhalation of 4-5 % isoflurane followed by cervical dislocation, if not otherwise stated.

### Induction of acute bacterial epididymitis in mice

Uropathogenic *Escherichia coli* (UPEC) strain CFT073 (characterized by (Welch et al., 2002) were provided by the Institute of Medical Microbiology, Justus-Liebig-University Giessen, Germany and cultured as described previously (Bhushan et al., 2008). To elicit an ascending canalicular infection, vasa deferentia were bilaterally ligated followed by an intravasal injection of UPEC (in sterile 0.9% NaCl) close to the cauda (5 µl containing 1 x 10^5^ CFU) using a Hamilton syringe. Control ‘sham’ mice underwent the same surgical procedure with an intravasal injection of 5 µl sterile 0.9% NaCl. Mice were sacrificed at day 1, 3, 5, 7, 10, 14 and 21 after infection by isoflurane narcosis and cervical dislocation. For each timepoint, 3-6 mice were used per experimental approach. For all subsequent approaches, at least two independent experiments were conducted.

### RNA extraction, RNASeq and whole transcriptome analysis

RNA was extracted from caput (segment 1-5, including the initial segment), corpus (segment 6-7) and cauda (segment 8-10) samples using QIAzol Lysis Reagent (Qiagen) following the manufactureŕs recommendation using a bead-based tissue homogenizer (Retsch, using 2.8 mm stainless steel beads). RNA was purified using the RNeasy Mini Kit (Qiagen) with an on-column DNase digestion using RNAse-free DNase Set (Qiagen) for 30 minutes to eliminate genomic DNA contamination. Total RNA and library integrity was verified with LabChip Gx Touch 24 (Perkin Elmer, MA, USA). Ten ng total RNA was used as template for SMARTer® Stranded Total RNA-Seq Kit – Pico Input Mammalian (Takara Bio.) following the manufactureŕs recommendation.

Sequencing was conducted on the NextSeq500 instrument (Illumina, CA, USA) using v2 chemistry with 1×75 bp single end setup. The resulting raw reads were assessed for quality, adapter content and duplication rates with FastQC (Andrews, 2010). Trimmomatic version 0.39 was employed in order to trim reads after a quality drop down below a mean of Q20 in a window of 5 nucleotides (GRCm38.p5) using STAR 2.6.1d with the parameter “—outFilterMismatchNoverLmax 0.1” to increase the maximum ratio of mismatches to mapped length to 10% (Dobin et al., 2013). The number of reads aligning to genes was counted with the featureCounts 1.6.5. tool from the Subread package (Liao et al., 2013). Only reads mapping at least partially inside exons were admitted and aggregated per gene. Reads overlapping multiple genes or aligning to multiple regions were excluded. Differentially expressed genes were identified using DESeq2 Version 1.18.1 (Love et al., 2014). Only genes with a minimum fold change of ±1.5 (log2 ± 0.59), a maximum Benjamini-Hochberg corrected p-value of 0.05, and a minimum combined mean of 5 reads were considered to be significantly differentially expressed. The Ensembl annotation was enriched with UniProt data (release 06.06.2014) based in Ensembl gene identifiers (UniProt Consortium, 2014.)

### Determination of colony forming units (CFU)

For each time point (1, 3, 5, 7 and 10 days post-infection), four biological replicates were used to assess the bacterial loads in the different epididymal regions. Data were obtained from two independent experiments. Tissue was collected and separated under sterile conditions in the IS, caput, corpus and cauda before homogenization in 250 µl sterile ice-cold PBS. Ten-fold series dilutions were prepared and spread onto Luria broth (LB) agar plates (10 mg/ml tryptone, 5 mg/ml yeast extract, 10 mg/ml NaCl and 15 mg/ml agar agar (pH 7.0)). Plates were incubated up-side-down at 37°C overnight before were colony forming units (CFU) were counted and calculated in relation to the previously determined tissue weight (per mg of used tissue). Pure *E.coli* were plated as positive control, whereas PBS only that was kept in similar tubes as the samples prior to plating to exclude contaminations within PBS solution and used tubes.

### Histological staining (modified Masson-Goldner trichrome staining)

Bouin’s-fixed (5 hours) and paraffin-embedded epididymides were cut into 5 µm sections. Deparaffinized and rehydrated tissues were stained for two minutes with Weigert’s iron hematoxylin for nuclear labeling (1:1 mixture of stock solution I (10 mg/ml hematoxylin 96% ethanol) and stock solution II (11.6 mg/ml FeCl_3_ in 2.5% HCl) followed by blueing in running tap water for 15 minutes. Sections were rinsed in 1% acetic acid followed by cytoplasmic staining with Ponceau-Acid Fuchsin (10 mg/ml Ponceau de Xylidine, 5 mg/ml Acid Fuchsin in 2% acetic acid) for 5 minutes. Sections were rinsed in 1% acetic acid for 3 minutes before incubated in 5% phosphotungstic acid for 30 minutes (under visual control). Sections were rinsed in distilled water for 3 minutes, followed by staining of connective tissue using aniline blue – orange G solution (5 mg/ml aniline blue and 20 mg/ml Orange G in 8% acetic acid for 30 minutes. Subsequently, sections were rinsed in 1% acetic acid followed by dehydration in increasing concentrations of ethanol and xylene and mounting using Entellan (Sigma-Aldrich). Images were acquired using a Leica DM750 microscope (Leica Microsystems, Wetzlar, Germany). In order to create whole organ images, single-captured images were composed using Inkscape V0.92.4. Morphometric analyses were performed using ImageJ V1.53a.

In order to assess the luminal diameter in different epididymal regions, 20-30 duct cross sections were measured per segment from the opposite apical surfaces of the ductal epithelium using ImageJ V1.53a. Measurements from segment 1-5 were averaged and summarized as ‘caput’ (incl. IS). Measurement from segment 8-10 were averaged and summarized as ‘cauda’. In total, three biological replicates were used per infection time point and experimental group.

### Disease score of acute bacterial epididymitis

A disease scoring system was slightly modified from a previously reported disease score established for experimental autoimmune epididymo-orchitis (Wijayarathna et al., 2020), in order to categorize and compare the observed histopathological alterations throughout the time course. The scoring system considered the following aspects:

0 – No histological alterations, normal tissue architecture
1 – Scattered/ focal mild histological alterations
2 – Mild histological alterations (mild reduction of the luminal diameter, mild interstitial fibrosis)
3 – Mild to moderate histological alterations (mild interstitial fibrosis, moderate luminal diameter reduction, focal and mild epithelial damage)
4 – Moderate histological alterations (moderate interstitial fibrosis, moderate luminal diameter reduction, moderate epithelial damage)
5 – Severe histological alterations (severe interstitial fibrosis, severe luminal diameter reduction, loss of epithelial integrity, presence of “ductal ghosts”)

### *Ex vivo* organ culture and cytokine measurement

Epididymides were isolated from 10-12 weeks old C57BL/6J mice and separated into IS (segment 1), caput (segment 2-5), corpus (segment 6-7) and cauda (Segment 8-10, Suppl. Fig. 1E, n=4). Organ pieces were transferred into a 24-well plate containing RPMI media only and then pre-incubated for 15 minutes at 34 °C with 5 % CO_2_ before 50 ng/ ml lipopolysaccharide (LPS) was added. After 6 hours incubation, supernatants and organ pieces were collected. Protein concentrations of inflammatory cytokines were determined by LegendPlex™ Multiplex Assay (BioLegend) using the pre-defined mouse inflammation panel according to the manufactureŕs instructions. Cytokine levels were determined in both tissue homogenate (protein extraction using RIPA-buffer and quantification using Bradford Assay) and the supernatant, producing similar results.

### Gentamicin assay

Epididymides were isolated from 10-12 weeks old C57BL/6J mice and cultured *ex vivo* as described above (see ‘*Ex vivo* organ culture and cytokine measurements’). 1 x 10^6^ uropathogenic *E. coli* were added to the organ culture and incubated for 4 h. Supernatants were carefully removed and organ pieces were washed twice with sterile PBS. Subsequently, organ pieces were incubated within 1 ml RPMI media containing 200 µg/ml gentamicin for 1 h at 34°C and 5% CO_2_ resulting in elimination of extracellular bacteria while intracellular bacteria were unaffected. Organ pieces were washed twice with PBS prior to tissue homogenization in 250 µl sterile ice-cold PBS. Homogenates were spread onto LB agar plates and incubated up-side-down 24 h at 37°C. Colonies were counted and calculated in relation to the previously determined tissue weight (per 10 mg of used tissue). Data were obtained from two independent experiments. UPEC alone were plated as positive control. A bacterial suspension that was treated with gentamicin under the same conditions as the organ pieces showed no colony forming as proof of antibiotic effectiveness.

### Cell preparation and surface staining for flow cytometry

For flow cytometric analyses, mice were sacrificed by deep isoflurane anesthesia and cervical dislocation. In order to eliminate the majority of intravascular CD45^+^ cells, mice were perfused with PBS by inserting a 30G needle into the left ventricle of the heart, while the right ventricle was opened by a small incision. Up to 50 ml PBS were carefully and continuously injected for 5-10 minutes until the tissue in the scrotal area cleared, (especially the highly vascularized initial segment). For flow cytometric analyses of UPEC-infected mice, epididymides were separated into caput (containing the initial segment) and cauda and single organs were used for single cell suspension (due to individual differences in immune responses). For flow cytometric analyses under physiological conditions, epididymides were dissected into IS, caput, corpus and cauda and the tissue of three mice were pooled due to the small tissue size (5-7 mg per organ piece) in order to obtain sufficient numbers of cells. Collected tissue was mechanically dissociated by chopping followed by enzymatic digestion for 45 minutes at 37°C in DMEM containing collagenase D (1.5 mg/ml, Roche) and DNase I (60 U/ml, Sigma). Digested suspensions were aspirated through 30G needles four to six times and filtered through a 70 µm cell strainer before centrifugation at 400 x *g* for 10 min at 4°C. Single cell suspensions were incubated with red blood cell lysis (RBC lysis buffer, Qiagen) for three minutes before pelleted by centrifugation (400 x *g*, 10 minutes at 4 °C). Cells were re-suspended in PBS and stained with a fixable viability dye in order to assess viability [different viability dyes were used depending on the respective panel: ZombieAqua^TM^ (Biolegend, 423101), Zombie NIR^TM^ (BioLegend, 423105), Viobility^TM^ 405/452 Fixable Dye (Miltenyi, 103-109-816)] following the respective manufactureŕs recommendation. To block unspecific binding of antibodies to mouse cells expressing Fc receptors, cell suspensions were incubated with Fc blocking reagent (Miltenyi, 130-092-575) following manufacturers recommendations. Cells were stained with antibodies listed in the Key Resource Table (Table 1) for 30 minutes at 4°C in 50 µl MACS Quant buffer (2 mM EDTA and 0.5 % BSA in PBS). Respective controls were used by omitting the target antibody and incubating with the respective isotype control under same conditions. Cells were washed twice with MACS Quant buffer before re-suspension in 200-500 µl MACS Quant buffer (depending on cell numbers). Flow cytometry was performed using a MACS Quant Analyzer 10 and analyzed with FlowJo software version 10.5.3.

**Table 1:**
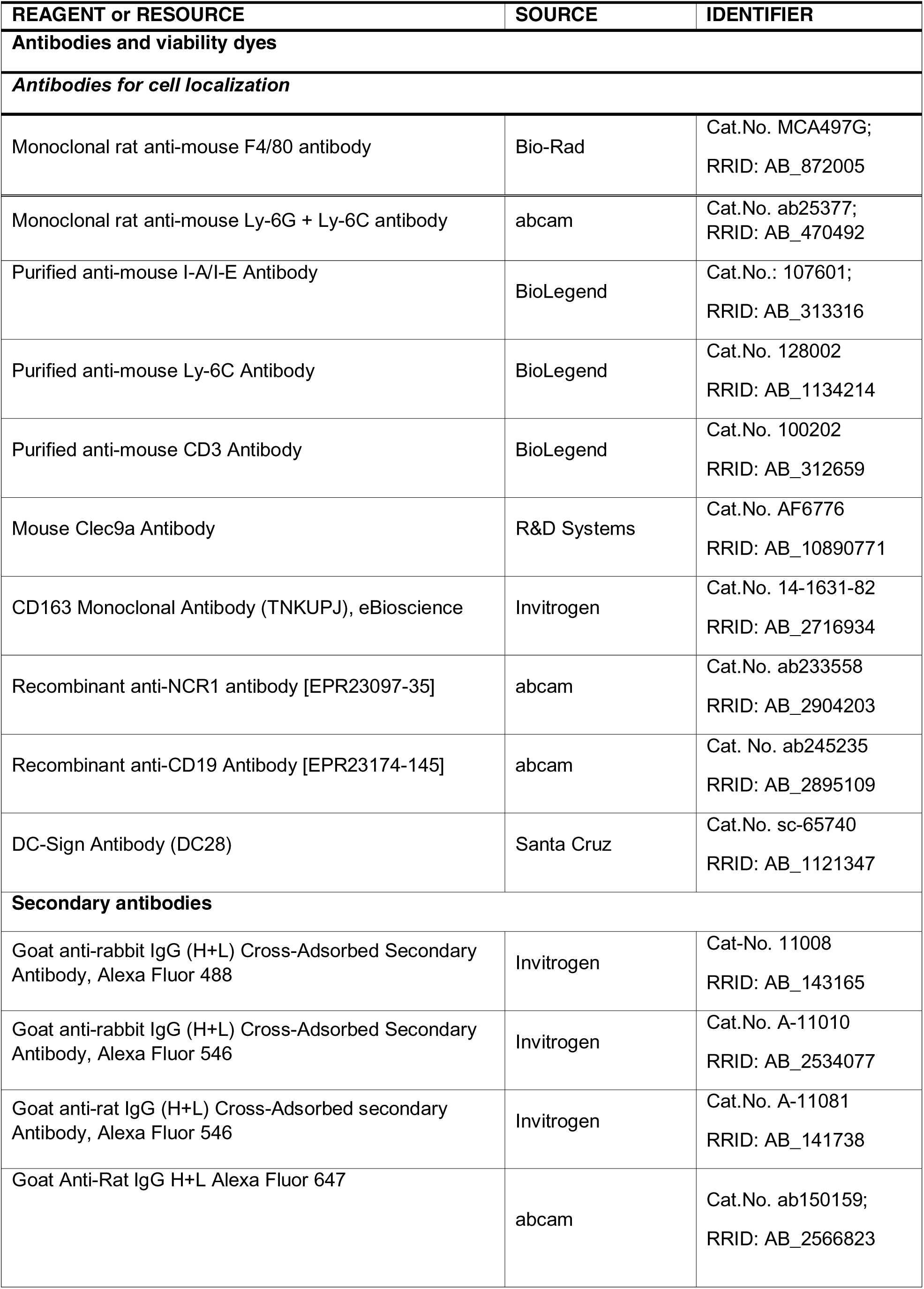
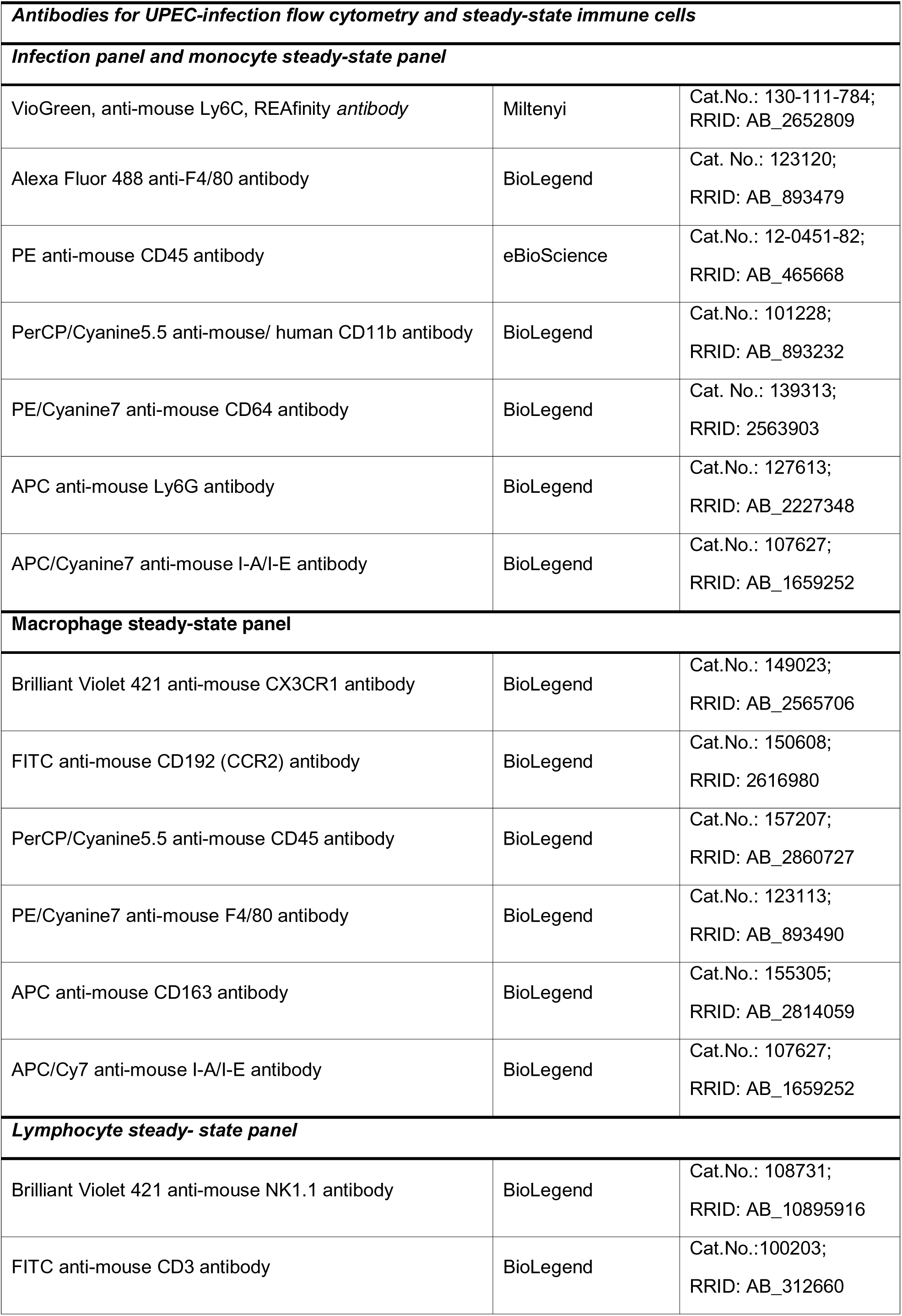
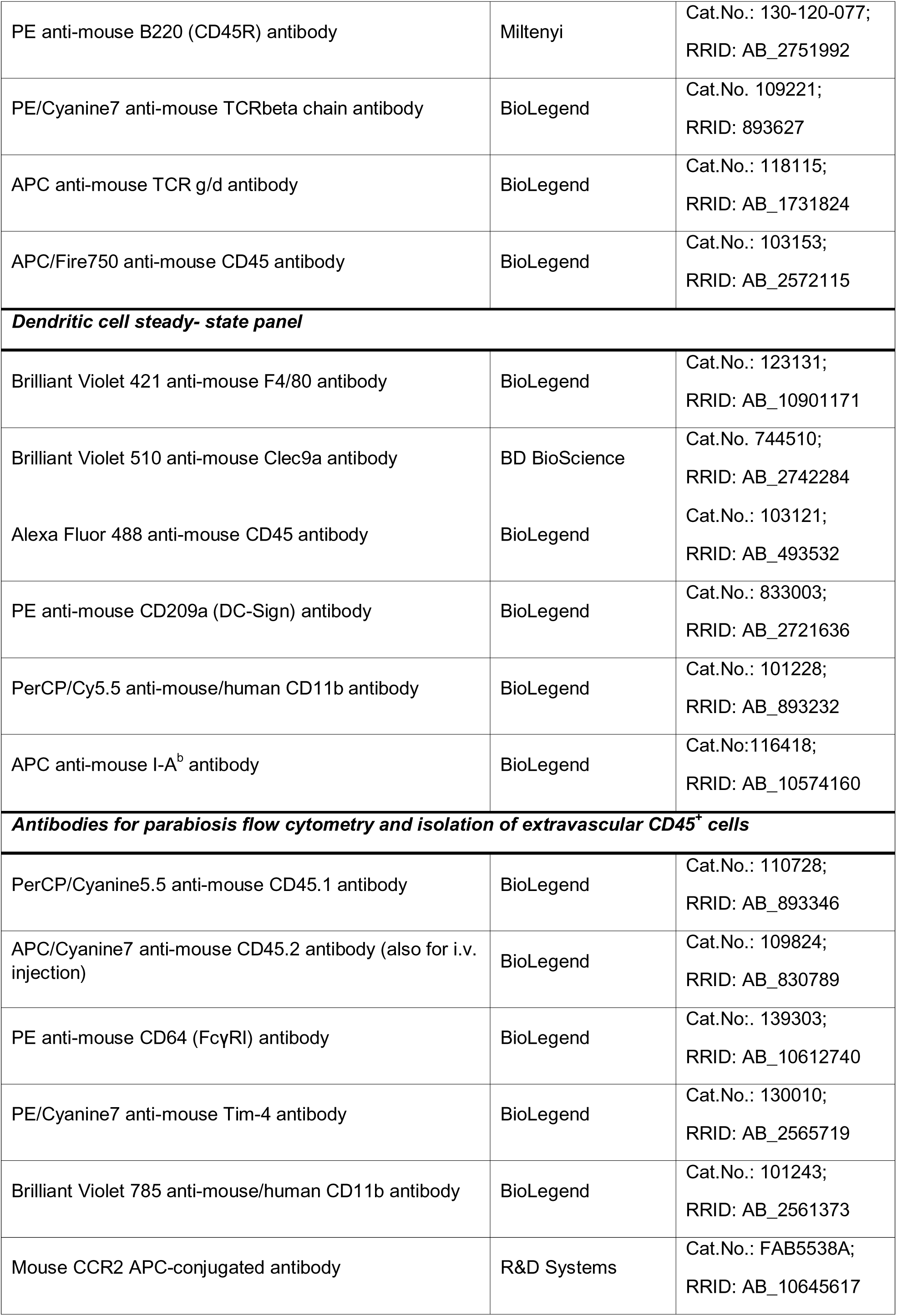
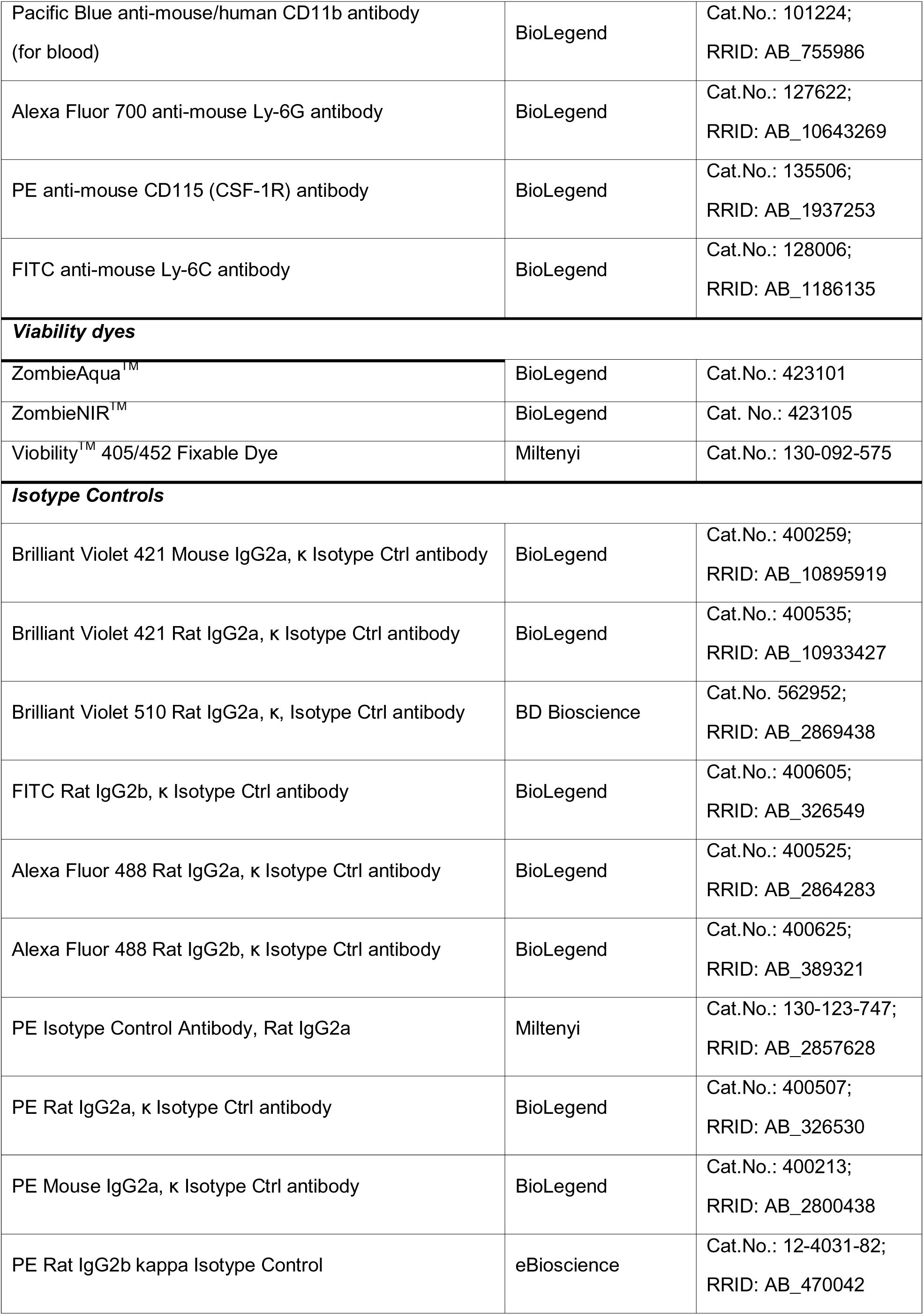
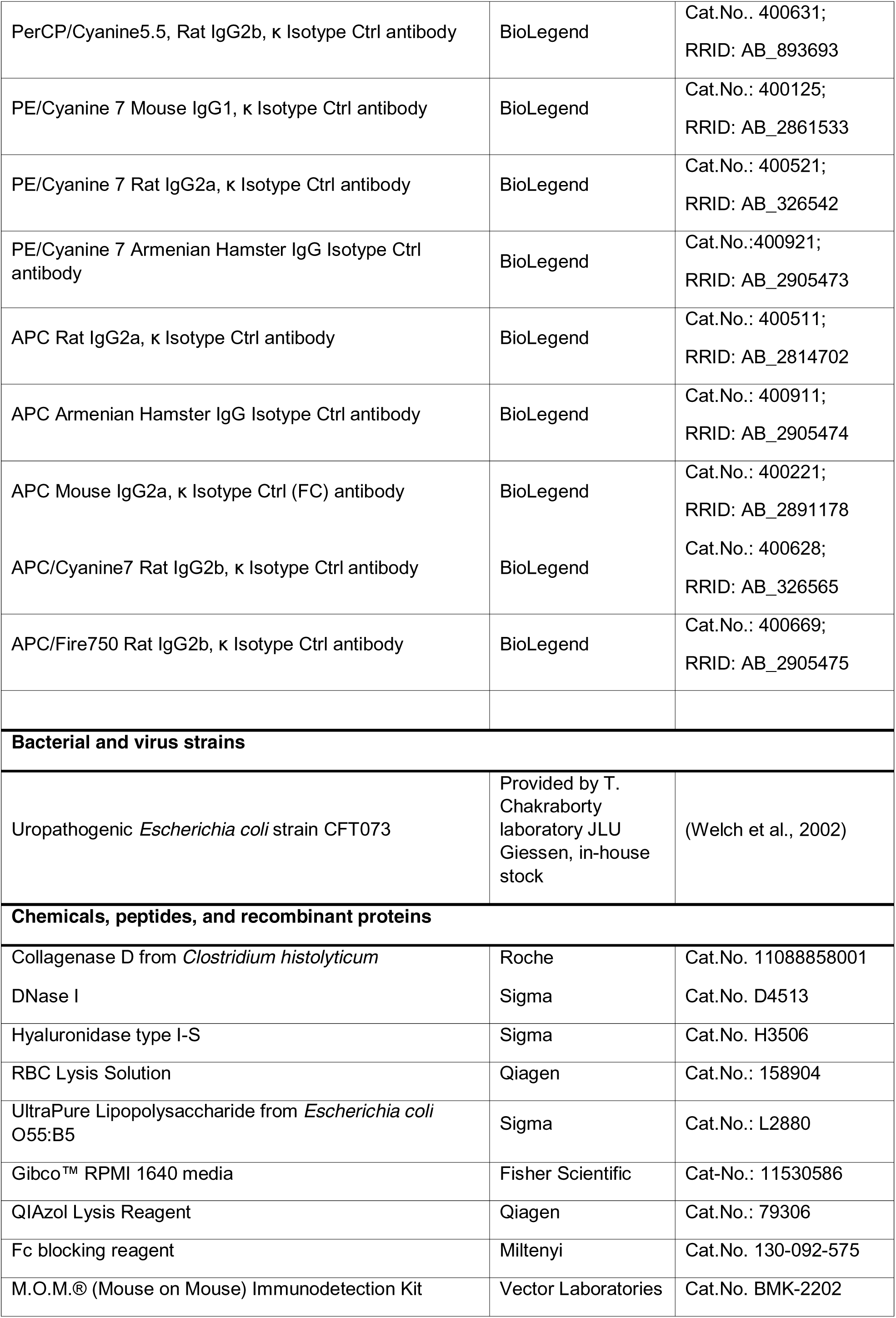
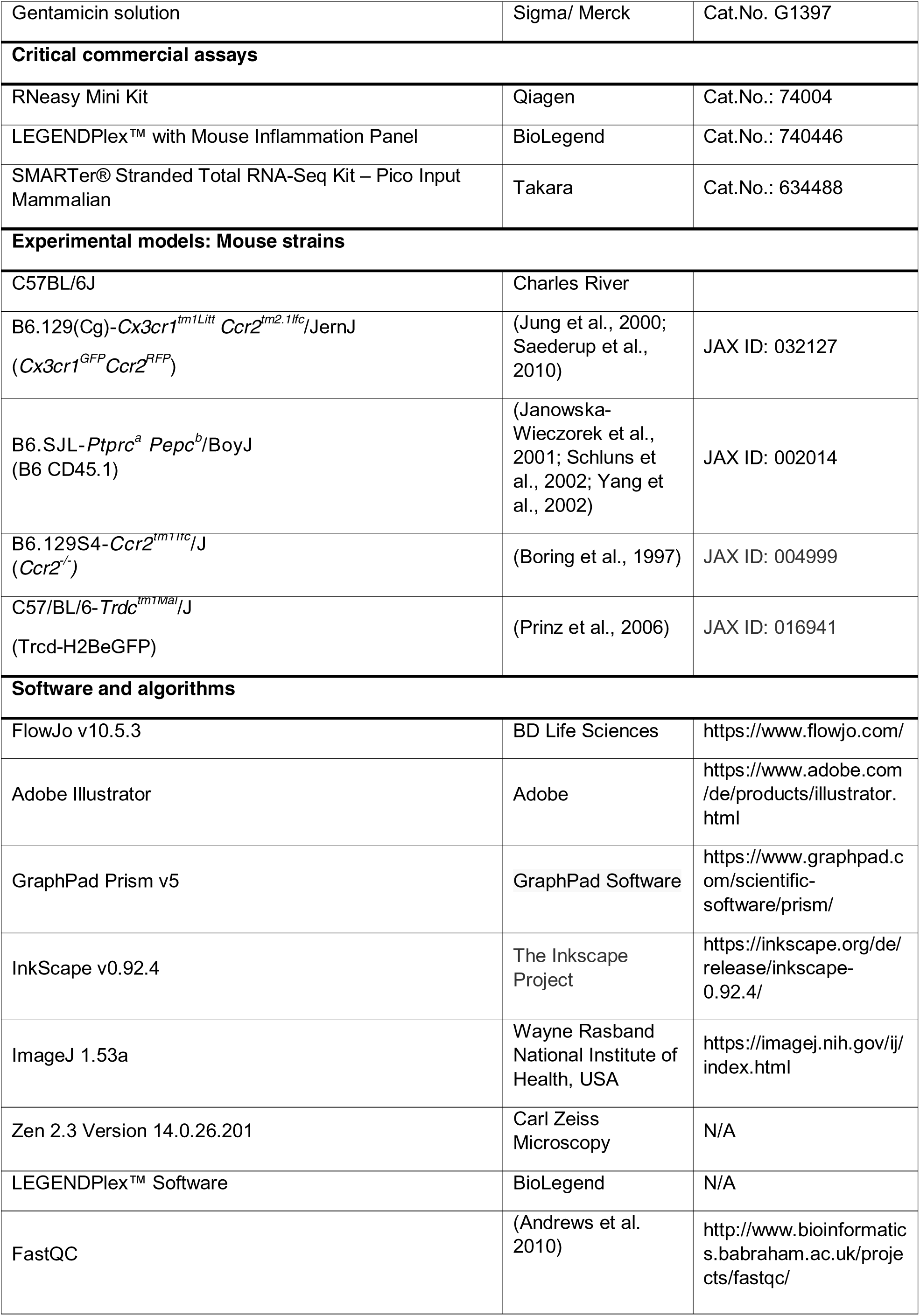
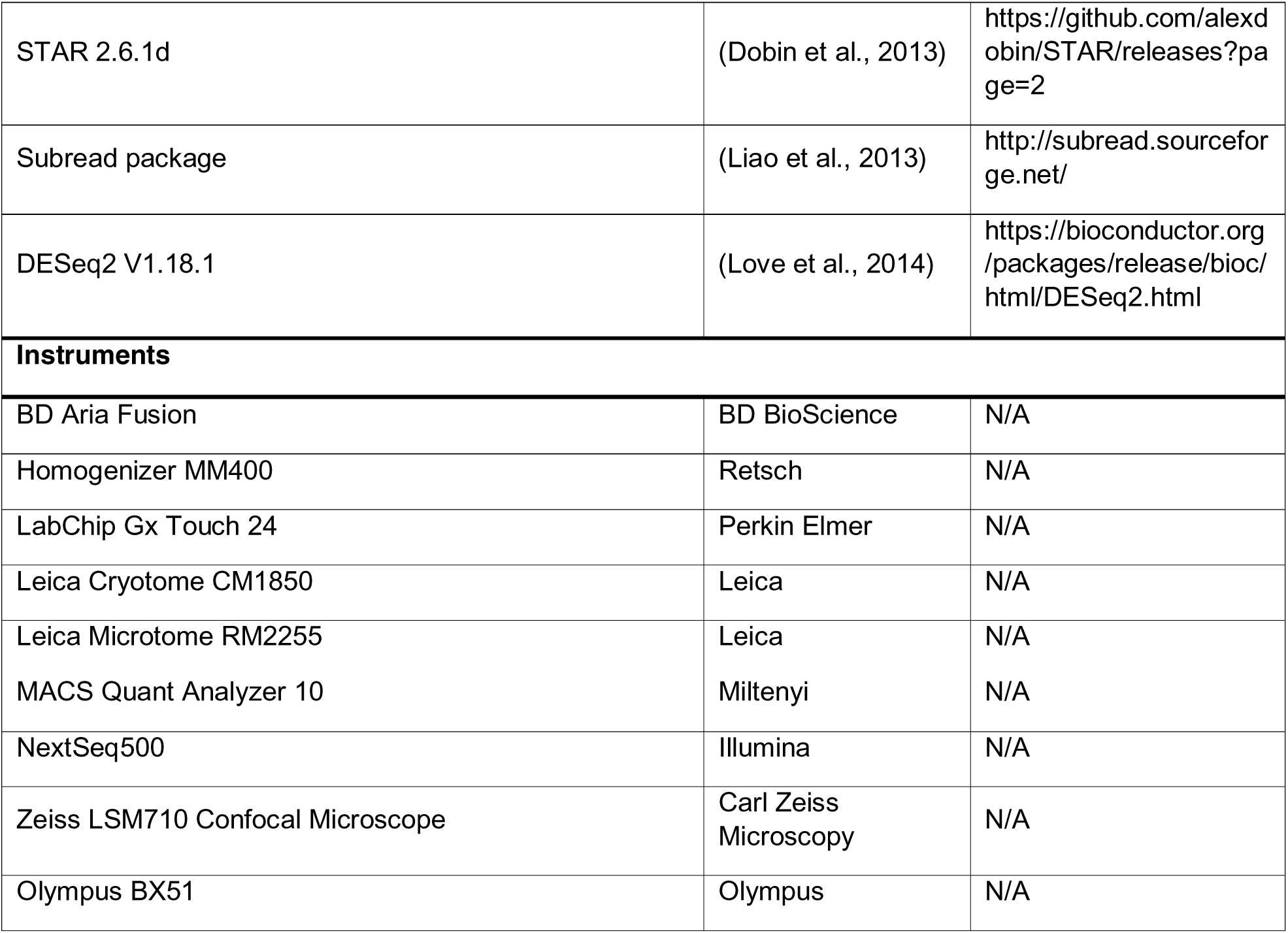
Key Resource Table.

### Gating strategy and panel constellation for flow cytometry

A general gating strategy (outlined in Suppl. Fig. S4A) was performed on each sample prior to the gating based on surface staining. Briefly, debris and sperm were excluded by SSC-A/ FSC-A followed by two-step single cell gating (FSC-H/FSC-A and SSC-H/SSC-A) and Live/Dead discrimination using viability dyes listed in the key resource table (table 1).

### Gating of immune cells in UPEC-infected and sham mice (Suppl. Fig. S1G) and monocytes in steady-state (Suppl. Fig. S4B)

#### Panel

Viobility^TM^ 405/452 Fixable Dye, CD45-PE, Ly6G-APC, Ly6C-VioGreen, CD11b-PerCP/Cy5.5, F4/80-AF488, CD11b-PerCP/Cy5.5, CD64-PE/Cy7, MHC-II-APC/Cy7 – see ‘Infection panel and monocyte steady-state panel’ in table 1 (key resource table) for more details.

*Neutrophils:*

CD45^+^Ly6G^+^
*Monocytes:*

CD45^+^Ly6G^-^Ly6C^+^CD11b^hi^
*Inflammatory monocyte-derived macrophages:*

CD45^+^Ly6G^-^Ly6C^+^CD11b^hi^F4/80^+^CD64^+^MHC-II^+^
*MHC-II^hi^ macrophages:*

CD45^+^Ly6G^-^Ly6C^-^CD11b^+^F4/80^hi^CD64^hi^MHC-II^+^
*MHC-II^lo^ macrophages:*

CD45^+^Ly6G^-^Ly6C^-^CD11b^+^F4/80^hi^CD64^hi^MHC-II^lo^
*Total DC (containing cDC1 and cDC2):*

CD45^+^Ly6G^-^Ly6C^-^ F4/80^-^CD64^-^MHC-II^hi^CD11b^lo-hi^

All CD45^+^ cells that were negative for all investigated markers (see above) were broadly considered to be lymphocytes.

*Monocytes steady-state:* CD45^+^F4/80^lo^CD11b^hi^Ly6C^hi^

### Gating of immune cells under physiological conditions

#### Dendritic cells

##### Panel

F4/80-BV421, Clec9a-BV510, CD45-AF488, CD209a-PE, CD11b-PerCP/Cy5.5, MHC-II-APC, ZombieNIR™ Fixable Dye – see ‘Dendritic cell steady-state panel’ in table 1 for more details.

*Conventional dendritic cells 1 (cDC1):* CD45^+^F4/80^-^MHC-II^hi^Clec9a^+^
*Conventional dendritic cells 2 (cDC2):* CD45^+^F4/80-MHC-II^hi^CD209a^+^

#### Lymphocytes

##### Panel

NK1.1-BV421, ZombieAqua™ Fixable Dye, CD3-FITC, B220 (CD45R)-PE, TCRbeta-PE/Cy7, TCRgd-APC, CD45-APC/Fire750 – see ‘Lymphocyte steady-state panel’ in table 1 for more details.

*B cells:* CD45^+^CD3^-^B220^+^
*T cells:* CD45^+^B220^-^CD3^+^ NK1.1^-^
*αβ T cells:* CD45^+^B220^-^CD3^+^ NK1.1^-^TCRbeta^+^TCRγδ^-^
*γδ T cells:* CD45^+^B220^-^CD3^+^ NK1.1^-^TCRbeta^-^TCRγδ^+^
*NK cells:* CD45^+^B220^-^CD3^-^NK1.1^+^

#### Macrophage subpopulation steady-state

##### Panel

CX3CR1-BV421, ZombieAqua™ Fixable Dye, CCR2-FITC, CD45-PerCP/Cy5.5, F4/80-PE/Cy7, CD163-APC, MHC-II-APC/Cy7 – see ‘Macrophages steady-state panel’ in table 1 for more details.

Cluster 1: F4/80^+^CX3CR1^hi^CCR2^-^MHC-II^-^
Cluster 2+6: F4/80^+^CX3CR1^hi^CCR2^-^MHC-II^+^
Cluster 3+4: F4/80^+^CX3CR1^+^CCR2^+^MHC-II^+^
Cluster 5: F4/80^+^CD163^+^CCR2^-^
Cluster 7: F4/80^+^CX3CR1^lo^CCR2^-^MHC-II^+^
Cluster 8: F4/80^+^CX3CR1^lo^CCR2^-^MHC-II^-^
Cluster 9: F4/80^+^CD163^+^CCR2^+^MHC-II^+^

### Parabiosis

Male donor mice (B6 CD45.1, JAX ID: 002014, (Janowska-Wieczorek et al., 2001; Schluns et al., 2002; Yang et al., 2002)) and recipient mice (CD45.2 *Ccr2^-/-^* JAX ID: 004999, (Boring et al., 1997)) were laterally shaved and conjoined by matching skin incisions from behind the ear to the tail as described previously (Dick et al., 2019). Six months after parabiosis surgery, mice were sacrificed by CO_2_ inhalation prior to blood and organ collection. In total, six recipient mice were analyzed. Cells were isolated from the four main epididymal regions (IS, caput, corpus, cauda) and epididymal fat for flow cytometry as outlined above. The chimerism for each macrophage sub population was normalized to blood monocytes in the recipient mouse (% normalized chimerism = (%donor cells in recipient/ %Ly6C^hi^ monocyte donor cells in recipient) *100) according to (Dick et al., 2019).

Gating strategy (as outlined in Suppl. Fig. S7A): Debris and sperm were excluded by SSC-A/FSC-A followed by a two-step doublet exclusion based on FSC-H/FSC-A and SSC-H/SSC-A. Total resident macrophages from all epididymal regions and epididymal fat were identified as CD45^+^CD11b^+^CD64^+^. TIMD4^+^ macrophages and CCR2^+^ macrophages were gated as internal controls for self-renewing and monocyte-derived macrophages, respectively (according to (Dick et al., 2022)). CD45^+^CD11b^+^CD64^+^CCR2^-^TimD4^-^ macrophages represented the entirety of all resident macrophage subpopulations. Blood monocytes were identified as CD45^+^Ly6G^-^CD115^+^CD11b^+^Ly6C^+^.

### Single cell preparation of extravascular CD45^+^ cells

Ten 10-week-old male wild type C57BL/6J mice were intravenously injected with an APC/Cyanine 7-conjugated anti-mouse CD45.2 antibody (Clone 104, BioLegend 109824, RRID: AB_830789) five minutes prior to euthanasia by CO_2_ inhalation. Single cell suspensions of epididymal regions (IS, caput, corpus, cauda) were prepared as previously described (see Cell preparation and surface staining for flow cytometry), with inclusion of 60 U/ml hyaluronidase type I-S (Sigma, H3506) in the digestion buffer. Cells were stained with a PerCP-Cyanine 5.5-conjugated CD45.1 antibody (BioLegend 110728, RRID: AB_893346). Stained single cell suspensions of all mice were pooled to obtain enough immune cells for sorting. Single live CD45.1^+^CD45.2^-^ immune cells were sorted on the BD Aria Fusion (BD Bioscience) for single cell RNA sequencing.

### Library preparation and data analysis

Single cell suspensions were prepared as outlined previously (Dick et al., 2019; Dick et al., 2022) using the 10x Genomics Single Cell 3’ v3 Reagent Kit user guide based on individually calculated cDNA concentrations. Briefly, cell suspension were washed twice with PBS supplemented with 0.04% BSA and centrifuged at 330 x g for 6 minutes. The appropriate volume for droplet generation was assessed by counting live cells using Tryptan Blue staining and a hemocytometer. Reverse transcription was performed in a pre-chilled 96-well plate (heat-sealed) using a Veriti 96-well thermal cycler (Thermo Fisher). cDNA was recovered using 10x-associated Recovery Agent followed by amplification and purification using SPRIselect beads (Beckman) following manufactureŕs recommendation. After diluting samples in a 4:1 ratio (elution buffer [Qiagen]:cDNA), cDNA concentration was determined using a Bioanalyzer (Agilent Technologies).

Sequencing libraries were produced by loading samples on the 10x Chromium. Generated libraries were processed as recommended by the methods provided from 10x Genomics. Expression matrices were generated using Cell Ranger (10x Genomics). Obtained raw base call (BCL) files from the HiSeq2500 sequencer were demultiplexed into FastQ files. Sequencing reads were aligned to the mouse genome/ transcriptome (mm10) and counted by StarSolo. After library preparation and cell mapping (StarSolo), 13015 data points were identified as valid cells (2076 within IS, 3791 within caput, 4523 within corpus, 2625 within cauda). Preprocessed counts were further analyzed using Scanpy. Basic cell quality control was conducted by taking the number of detected genes and mitochondrial content into consideration. In total, 49 cells, that did not express as least 300 genes or had a mitochondrial content greater than 10%, were removed. Genes were filtered out if they were detected in less than 30 cells (<0.2%). Raw counts per cell were normalized to the median count over all cells and transformed into log space to stabilize variance. Dimensionality reduction was performed by Principal Component Analysis (PCA), retaining 50 principal components. Subsequent steps, e.g. low-dimensional UMAP embedding and cell clustering via community detection, were based on the initial PCA. Final data visualization was performed using Scanpy and CellxGene package.

### Immunofluorescence

Epididymides from *Cx3Cr1*^GFP^*Ccr2*^RFP^ reporter mice (JAX ID: 032127, (Jung et al., 2000; Saederup et al., 2010)), Tcrd-H2BEGFP, (JAX ID: 016941, (Prinz et al., 2006)) and C57/BL6J mice (Charles River) were fixed with ROTI®Histofix 4% (Carl Roth, Germany) for 5 hours followed by washing in phosphate buffer and incubation in 30% sucrose overnight at 4°C before embedding in OCT media and storage at -80°C. 20 µm cryo-sections were prepared using a Leica Cryotome CM1850 and air-dried for 20 minutes followed by a 20 minute post-fixation in 100 % methanol at -20°C. After washing in TBS-T (TBS + 0.05% Tween, pH 7.6), sections were permeabilized using 0.2% Triton-X-100 in TBS-T for 20 minutes at room temperature. Washed sections were incubated for 30 minutes in blocking solution (3% BSA, 10% normal goat serum in TBS-T). Primary antibodies (for further specification see key resource table, table 1) were diluted in blocking solution (F4/80 [Bio-Rad]: 10 µg/ml, Ly6G [abcam]: 1 µg/ml, MHC-II [BioLegend]: 5 µg/ml), CD163 [Invitrogen]: 5 µg/ml, LY6C [BioLegend]: 5 µg/ml, CD3 [BioLegend]: 10 µg/ml, Clec9a [R&D Systems]: 15 µg/ml, NCR1 [abcam]: 7 µg/ml, CD19 [abcam]: 8 µg/ml, DC-Sign [Santa-Cruz]: 10 µg/ml) and incubated over night at 4°C. Secondary antibodies were diluted in TBS-T according to manufactureŕs recommendation and incubated one hour in a dark chamber at room temperature. Sections were thoroughly washed four-times for 10 minutes in TBS-T before mounting with Invitrogen™ ProLong™ Gold Antifade Mountant with DAPI (ThermoFisher). Sections were imaged with a Zeiss LSM 710 confocal microscope and Zen Software Version 14.0.26.201.

