## Supplemental Figures for "The regional distribution of resident immune cells shapes distinct immunological environments along the murine epididymis"

**Supplemental Figure S1**

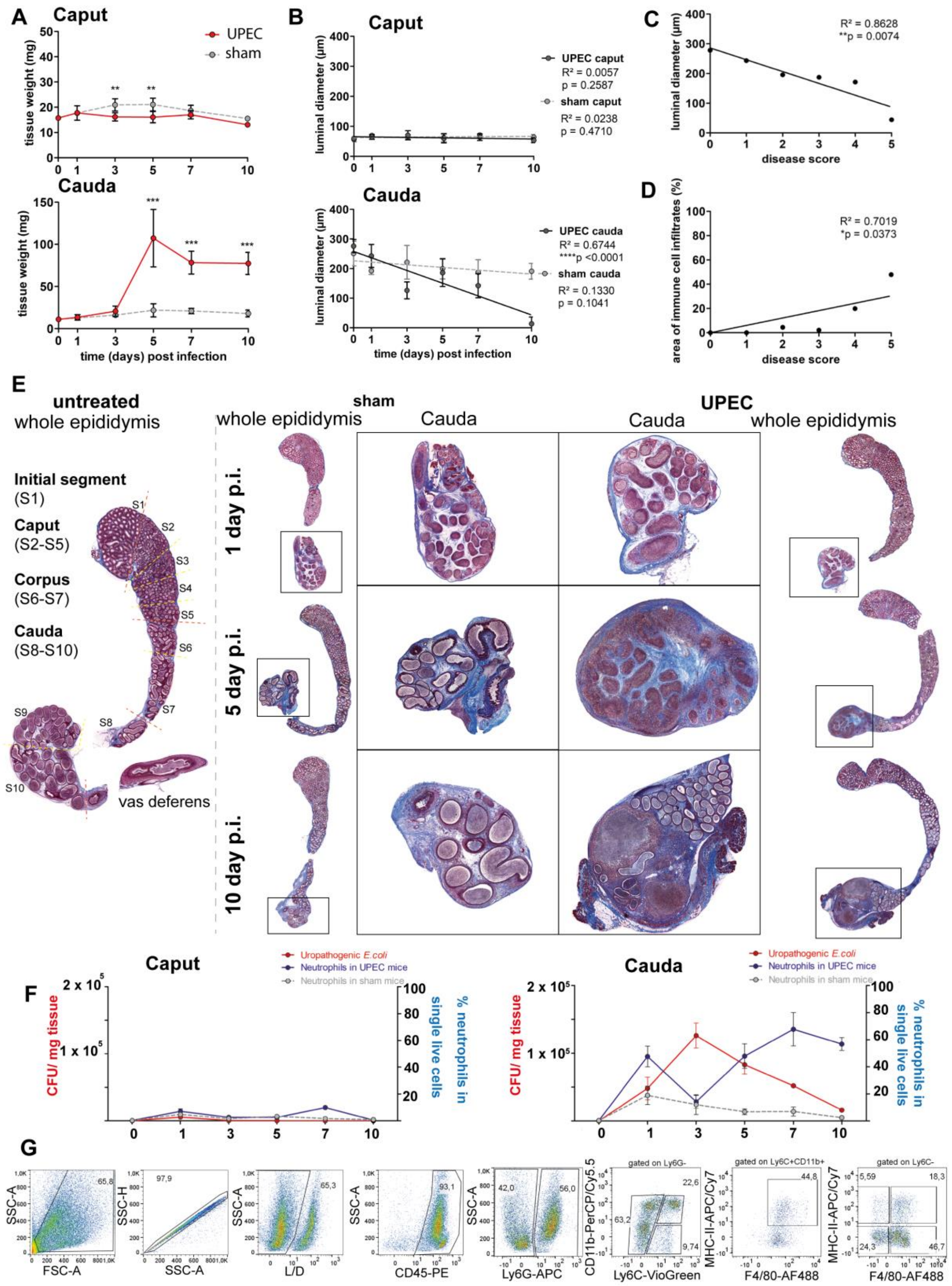


**Supplemental Figure 1: Morphometric analyses of the sham and UPEC-infected C57BL/6J wild type mice. Related to Figure 1 and 2.**

**(A)** Changes in weight (in mg) of caput and cauda epididymides throughout different infection time points post infection (mean ± SD, n=4-5, Two-way ANOVA with Bonferroni post hoc test, *p<0.05, **p<0.005, ***p<0.001).

**(B)** Luminal diameter of the epididymal duct (in µm) within caput and cauda epididymides. 30-50 duct cross sections were measured per region of the biological replicate and averaged (mean ± SD, n=4-5, Two-way ANOVA with Bonferroni post hoc test, *p<0.05, **p<0.005, ***p<0.001).

**(C)** Pearson correlation plot of disease score and luminal diameter. The average luminal diameter per disease score is shown. A Pearson´s correlation was considered to be statistically significant at p<0.05.

**(D)** Pearson correlation plot of disease score and area of immune cell infiltrates assessed by histological measurement. The average area of immune cell infiltrates per disease score is shown. A Pearson´s correlation was considered to be statistically significant at p<0.05.

**(E)** Histological images (modified Masson-Goldner trichrome staining) of the epididymis of untreated, sham and UPEC-infected C57BL/6J mice at different time points post infection.

**(F)** Percentage of infiltrating neutrophil granulocytes (blue) in relation to CFU/ mg tissue in caput and cauda epididymides of sham and UPEC-infected mice.

**(G)** Representative plots showing the gating strategy for flow cytometric analyses of infiltrating immune cell populations in sham and UPEC-infected mice.

**Supplemental Figure S2**

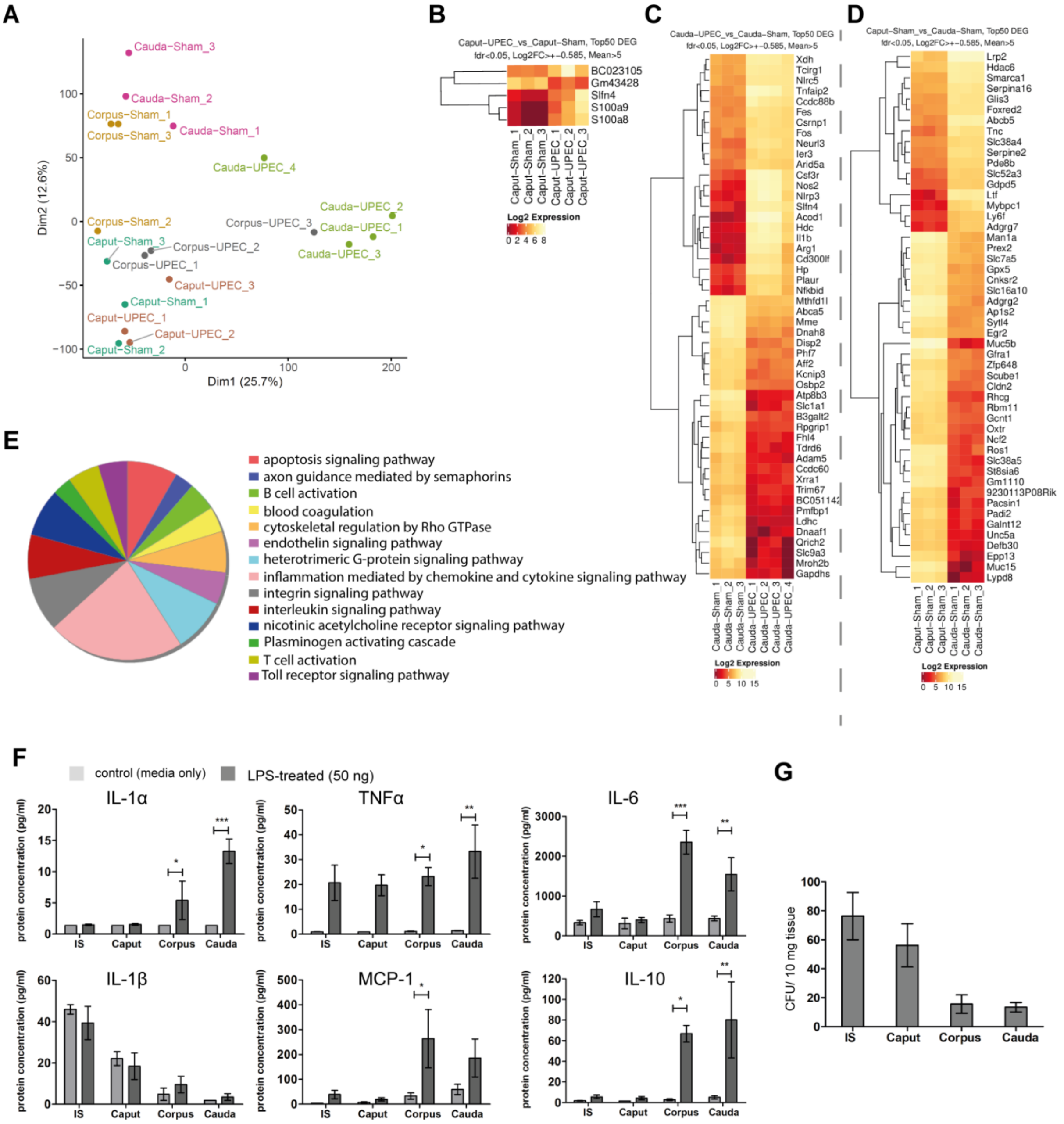

**Supplemental Figure 2: Differential immune responses within the epididymal regions in C57BL/6J mice assessed by RNASeq analyses of *in vivo* epididymitis and Multiplex assay-based determination of cytokine levels from *ex vivo* organ culture. Related to Figure 1.**

**(A)** Principal component analysis of all investigated *in vivo* epididymitis samples in RNASeq analysis (n=3-4 per goup).

**(B)** Heatmap showing DEG between the caput of sham and UPEC-infected mice *in vivo* – related to the Volcano plot shown in Fig. 1G, Cut-off criteria are indicated above the heatmap.

**(C)** Heatmap showing DEG between the cauda of sham and UPEC-infected mice *in vivo* – related to the Volcano plot shown in Fig. 1G, Cut-off criteria are indicated above the heatmap.

**(D)** Heatmap showing DEG between the caput sham and cauda of sham mice *in vivo* – related to the Volcano. Cut-off criteria are indicated above the heatmap.

**(E)** Pie chart showing upregulated gene sets and pathway enrichment within cauda epididymidis of UPEC-infected mice 10 days p.i. (compared to sham control mice, based on Panther database analysis).

**(F)** Indicated cytokines (IL-1α, IL-1β, TNFα, MCP-1, IL-6, IL-10) were measured within the culture media after *ex vivo* stimulation of the indicated epididymal regions with LPS (50 ng) for 6 hours.

**(G)** Bacterial uptake potential of the different epididymal regions (IS, caput, corpus, cauda) was determined by assessing the intracellular bacterial load after 4 h *ex vivo* organ culture with 1 x $10^6$ UPEC and subsequent treatment with gentamicin to eliminate extracellular bacteria (n=4, mean±SD).

### Supplemental Figure S3

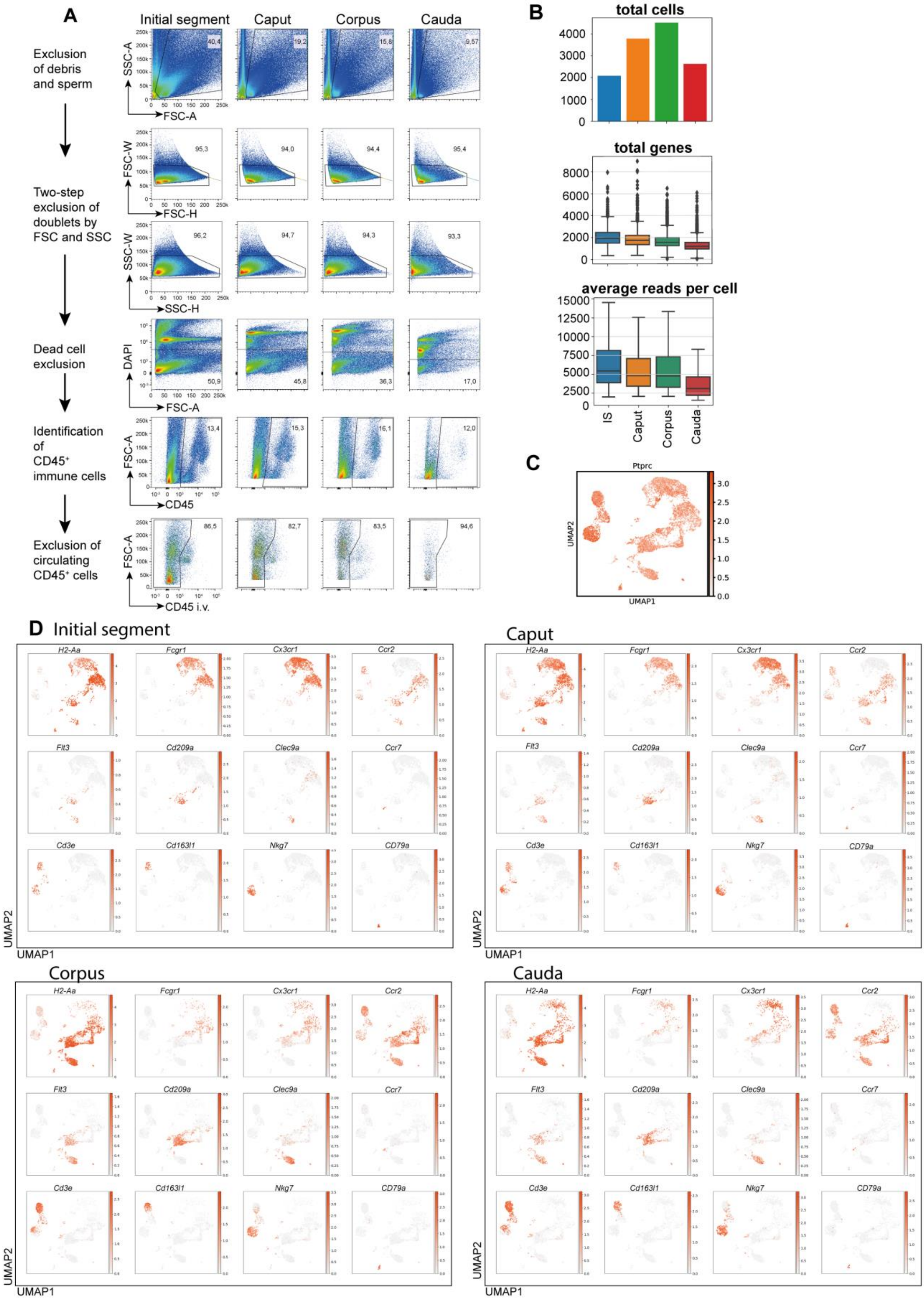


**Supplemental Figure 3: Isolation of extravascular CD45$^+$ cells for scRNASeq, quality controls for single cell reads and re-confirmation of identified CD45$^+$ populations by flow cytometry. Related to figure 3.**

**(A)** Extravascular CD45$^+$ cells of different epididymal regions were sorted following the indicated gating strategy prior to single cell RNASeq.

**(B)** Number of total cells, total genes and average reads per cells are indicated for different epididmal regions that were separately isolated.

**(C)** UMAP plot showing the expression of *Ptprc* (encoding CD45) in the identified cluster.

**(D)** UMAP plots showing the expression of selected key marker for the indicated immune cell population within the epididymal regions Initial segment, caput, corpus, cauda.

**Supplemental Figure S4**

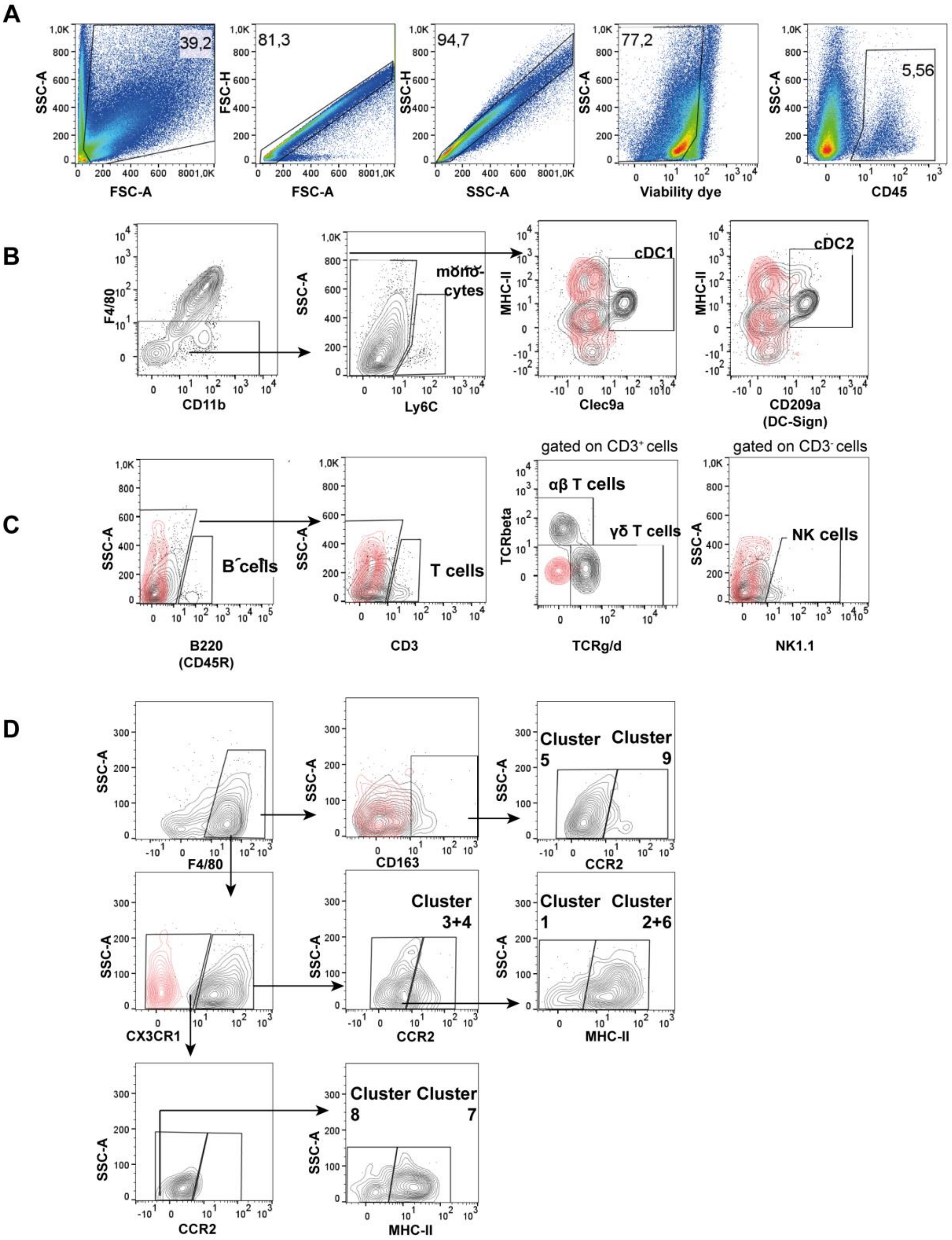


**Supplemental Figure 4: Gating strategy behind flow cytometry analyses of all immune cell populations under physiological conditions. Related to Fig. 5, Suppl. Fig. S3 and Suppl. Fig. S5.**

(**A**) General gating that has been applied to each sample included exclusion of debris und sperm based on SSC-A vs. FSC-A, followed by a two-step single cell gating (based on FSC and SSC), live cells were discriminated using a viability dye (see table 1) according to the respective panel all leukocytes were identified by CD45 staining.

**(B)** F4/80$^{-}$CD11b$^{lo\text{-}hi}$ cells were further distinguished by Ly6C to identify monocytes (Ly6C$^{+}$), and Ly6C$^{-}$ cells were segregated using MHC-II in combination with Clec9a and CD209a to differentiate cDC1 (MHC-$^{II}$hiClec9a$^{+}$) and cDC2 (MHC-II$^{hi}$CD209a$^{+}$), respectively.

**(C)** Lymphocytes were segregated into B cells (B220$^{+}$) and T cells (CD3$^{+}$) that were further differentiated into αβ and γδT cells, as well as NK cells (NK1.1$^{+}$ cells).

**(D)** Gating strategy of macrophage subsets according to obtained scRNASeq data. Arrows indicate the gating strategy and identified subsets.

**(A-D)** Representative plots are from the cauda due to the most diverse immune cell distribution in this region. Red overlays represent the respective isotype controls.

**Supplemental Figure S5**

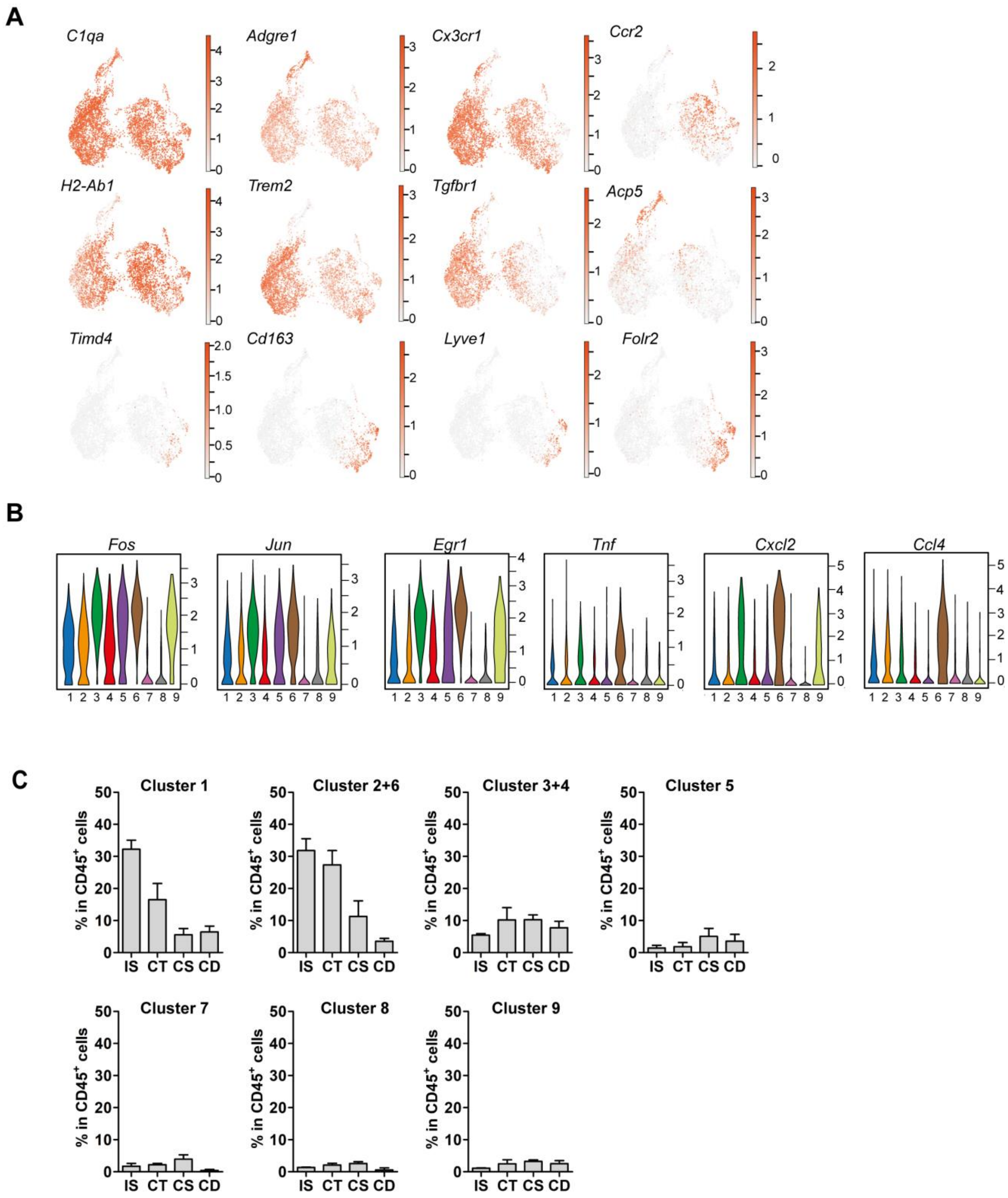


**Supplemental Figure 5: Distinct macrophage subgroups exist within the murine epididymis. Related to Fig. 4 and Fig. 5.**

**(A)** UMAP plots showing the expression of selected key marker for identified macrophage subgroups, related to Violin plots in Fig. 4D.

(**B**) Violin plots showing the expression of immediate-early activation genes (*Fos, Jun, Egr1*) as well as upregulated cytokines *Tnf, Cxcl2, Ccl4* among identified macrophage subgroups.

**(C)** Bar diagrams showing the percentage of identified macrophage subgroups within the CD45$^+$ population throughout the epididymal regions, assessed by flow cytometry and mirroring the distribution obtained by scRNASeq (n=4, mean ± SD).

### Supplemental Figure S6

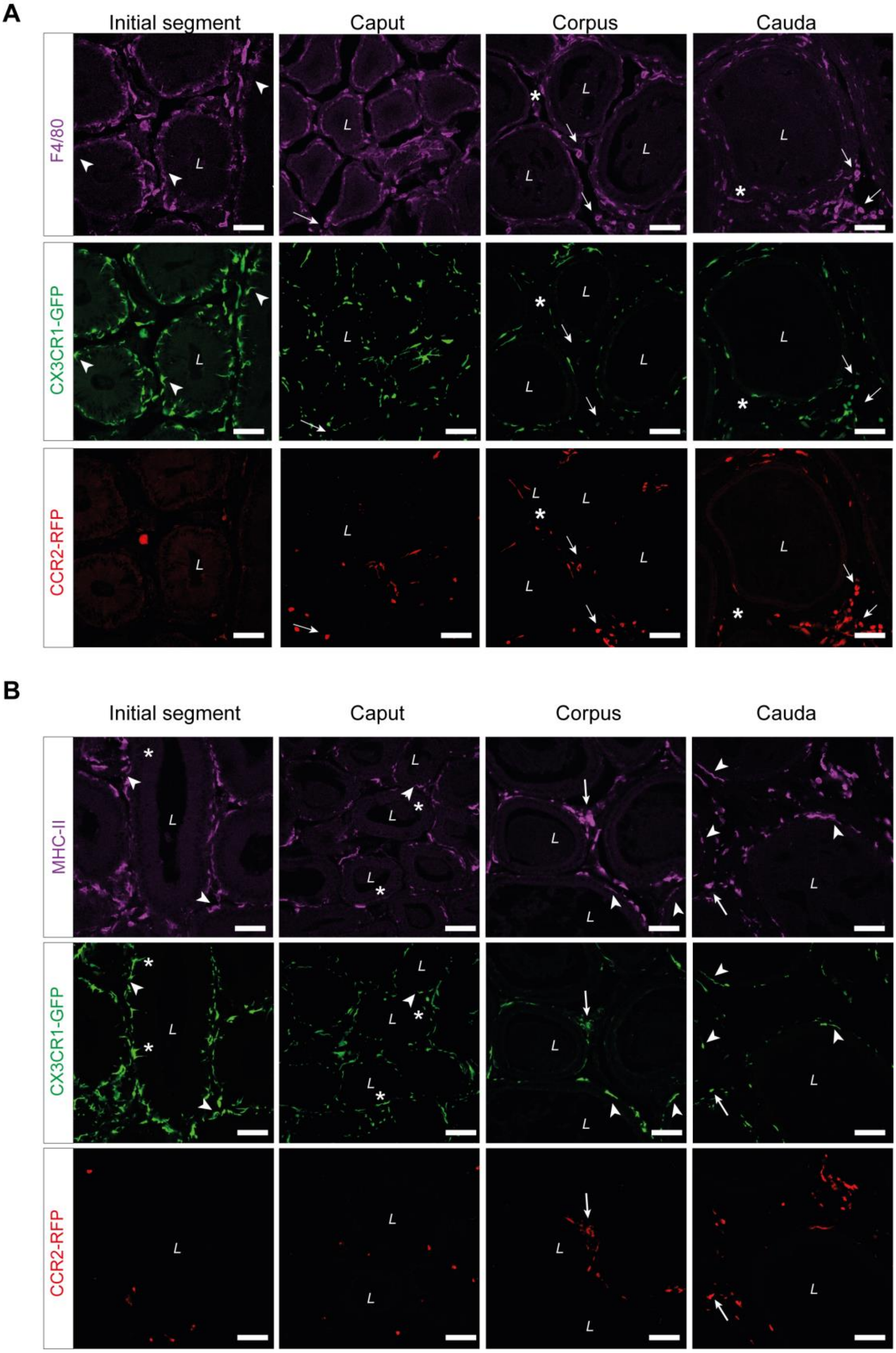


**Supplemental Figure 6: Confocal microscopy images showing the localization of identified macrophage subsets. Related to Fig. 5.**

(**A**) Single channel reads of anti-F4/80 (purple) staining on epididymal cryo-sections from adult *Cx3cr1*$^{GFP}$*Ccr2*$^{RFP}$ reporter mice. The majority of CX3CR1$^{+}$ cells were F4/80$^{+.}$ Arrowheads indicate the small fraction of intraepithelial F4/80$^{-}$ CX3CR1$^{+}$ cells within the IS. Arrows indicate interstitial F4/80$^{+}$ CX3CR1$^{-}$ cells that were CCR2$^{+}$ within caput, corpus and cauda epididymides. Asterisks (*) label a small fraction of F4/80 single positive cells ($CX3CR1^-CCR2^-$) found in the corpus and cauda. Scale bar 50 µm (L = Lumen).

**(B)** Single channel reads of anti-MHC-II (purple) staining on epididymal cryo-sections from adult *Cx3cr1*$^{GFP}$*Ccr2*$^{RFP}$ reporter mice. Asterisks (*) indicate intraepithelial $CX3CR1^+MHC\text{-}II^-$ cells within the IS and caput epididymides. Arrowheads indicate $CX3CR1^+MHC\text{-}II^+$ cells, lining the epididymal duct within the IS and situated within the epithelium within caput, corpus and cauda epididymides. Arrows indicate interstitial $CX3CR1^+MHC\text{-}II^+CCR2^+$ cells within corpus and cauda epididymides. Scale bar 50 µm (L = Lumen).

**Supplemental Figure S7**

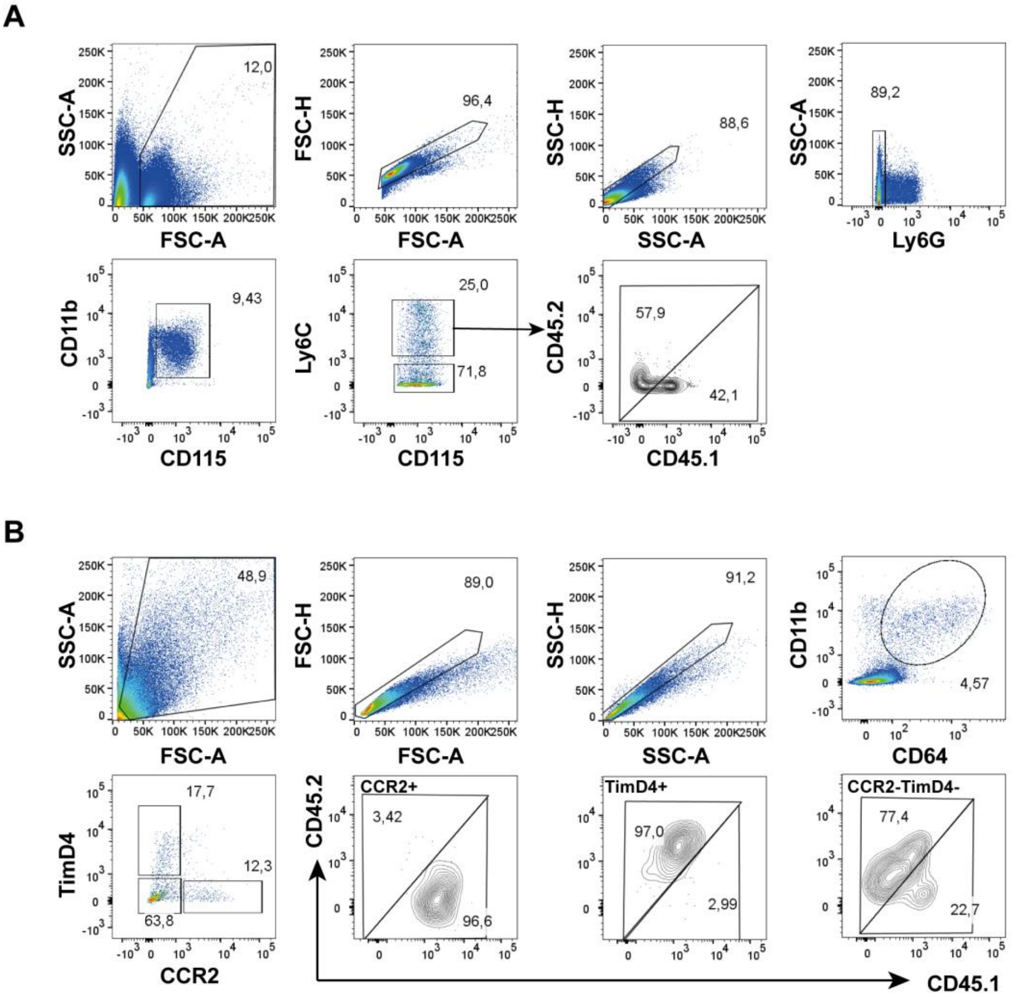


**Supplemental Figure 7: Gating strategy behind flow cytometric analyses of parabiosis experiments. Related to Fig. 6.**

**(A)** Gating strategy that was applied on blood samples from recipient *Ccr2*$^{-/-}$ mice. General gating was initially performed based on FSC and SSC in order to exclude debris and doublets, before neutrophils were excluded by selecting Ly6G$^{-}$ cells. Monocytes were further gated using CD115 and CD11b. Ratios of CD45.1$^{+}$ and CD45.2$^{+}$ events were assessed on Ly6C$^{+}$ monocytes.

(B) Representative plots (cauda) that were applied for the epididymal regions starting with general gating based on FSC and SSC in order to exclude debris and doublets. Total CD64$^{+}$CD11b$^{+}$ cells were gated and further segregated using TIMD4 and CCR2. The ratio of CD45.1 and CD45-2+ events was assessed in CCR2$^{+}$, TIMD4$^{+}$ and CCR2$^{-}$ TIMD4$^{-}$ cells.
